# Coordinated Division of Selective Stationary-Phase Yeast Cells Expands the Population Survivorship in Quiescence

**DOI:** 10.1101/2024.10.17.618937

**Authors:** Kai-Ching Hsiao, Hsin-Ying Lin, Tony R. Hazbun, Min-Hao Kuo

## Abstract

Budding yeast employs a variety of survival strategies in response to starvation, including pseudohyphal development, invasive growth, and sporulation. Here we report an innate pathway of “viability resurgence in quiescent time” (VRQT) by aged cultures to preserve and expand the population survivorship. Without additional nutrients, a few stationary-phase cells synchronously enter mitosis, increasing the population viability that does not likely result from mutations. VRQT is a population density-dependent collective behavior that can be triggered by quorum sensing aromatic alcohols. Genetic analyses demonstrate that VRQT is independent of many canonical pathways for cell growth, development, or stress responses. This community survival program allows yeast to proactively extend vitality amidst a common nutritional crisis. Wild strains isolated from clinical samples exhibit VRQT, suggesting that this survival mechanism must be considered when treating human infections.

## Introduction

Chronological lifespan of the budding yeast is a model for the survival of post-mitotic cells in metazoans. Typical laboratory strains have peak colony forming units (CFU) of about 3 - 5 x 10^8^ per mL that is equivalent to approximately 10 O.D._600_ absorption units of cells with close to 100% viability. In standard laboratory batch cultures, yeast cells go through a lag phase before engaging in logarithmic and fermentative growth until glucose is consumed. Diauxic shift follows and the cellular metabolism changes to utilize non-fermentable carbon sources in the environment. Cells enter the respiratory stationary phase when external carbon sources are exhausted(Gray et al., 2004). In the stationary phase, starvation drives yeast cells to a quiescent state where they temporarily exit the division cycle, but maintain the mitotic potential that supports propagation when resupplied with nutrients. As opposed to active and continual proliferation of cells in a typical laboratory batch culture, quiescence is the predominant state for all cells in their native environments(Sun and Gresham, 2021, O’Farrell, 2011). Prolonged starvation is thought to further cause quiescent cells to transition irreversibly into senescence, a metabolically active but mitotically deactivated state(Gray et al., 2004, Werner-Washburne et al., 2012). Yeast senescence is likened to the terminally differentiated human cells. These cells die eventually in this barren condition. Yeast cell death operationally refers to the loss of mitotic capacity, that is, the inability to divide a sufficient number of cycles to form visible colonies on fresh medium.

Interestingly, there are natural exceptions from the view of an irreversible and unstoppable journey to demise of cells. For example, genetic lineage tracing revealed that some mouse embryonic senescent cells lose their canonical senescence markers, re-enter the cell cycle, and continue to proliferate after birth, therefore challenging the notion of senescence being an irreversible state(Li et al., 2018). Two *S. cerevisiae* strains were isolated from a two-year old, sealed beer bottle(Aouizerat et al., 2019). Each of these long-lived strains possessed 30,000 nucleotide variations in the genome, with enrichment of genes associated with the target of rapamycin (TOR) pathway, DNA repair, mitochondrial functions, etc. Long-term survival of these strains therefore was suggested to result from mutations in the corresponding cellular functions. However, if these mutations indeed played a causal role in allowing cells to survive prolonged starvation, it is unknown whether such mutations arose prior to or during the biennial storage, or which mutations among the many varians that allowed survival. An elusive “adaptive regrowth” mutation(s) was thought to be responsible for the increase of viability (i.e., CFU) of stationary-phase cultures of laboratory and wild strains grown in synthetic complete medium(Fabrizio et al., 2004). This population revival was believed to depend on an altruistic programmed cell death mechanism(Fabrizio et al., 2004), a conclusion derived from the discoveries of mammalian apoptosis pathway of cell death in yeast(Fröhlich and Madeo, 2000, Madeo et al., 1997). A ∼1,500 Da cellular molecule was thought to be released from dying cells that transiently elevated the population viability(Herker et al., 2004). Altruistic or not, there is no evidence presented to show that the elevated CFU was due to cell division. The purported ∼1,500 Da cell-reviving molecule remains enigmatic.

The notion that acquired mutations lead to yeast adaptive regrowth(Fabrizio et al., 2004) or long-term survival(Aouizerat et al., 2019) follows the precedent of the bacterial GASP phenomenon(Finkel, 2006). GASP (growth advantage in stationary phase) is characterized by the capacity of cells to proliferate in long-term batch cultures without nutrient replenishment.

These cells possessed GASP mutations that not only caused aged cells to propagate in the depleted environment, but also, importantly, to outcompete younger cells from fresher cultures when grown together(Zambrano et al., 1993). However, the patterns of the GASP mutants outcompeting the young cells were far from uniform(Finkel, 2006). Some of the well-documented GASP mutations in *E. coli* change the alternative sigma (σ) factor RpoS or σ^s^ for expression of many stress responsive genes(Zambrano et al., 1993), and others affect high-affinity transporters for certain amino acids(Zinser and Kolter, 1999). GASP mutants appear to be short-lived in the aged population, because new mutants arise in waves, overtaking the previously established GASP population(Finkel, 2006). It is worth noting that the existence of adaptive regrowth mutations, or the yeast equivalent of GASP mutations, has not yet been proven. If they do exist, it is unclear why easily-acquired mutations that confer growth advantages have not been positively selected in the wild yeast strains, which are often threatened with starvation.

Aside from permanently changing their genetic makeup through spontaneous mutations, microorganisms use other means to survive stress. Many organisms engage in collective behaviors that enable groups of the members to coordinate their actions to overcome challenges that would be insurmountable for individuals(Couzin, 2009). Well-known examples of such behaviors include synchronous fireflies, insect swarms, fish schools, bird flocks, slime molds, and bacterial swarming and development(Reid and Latty, 2016). Synchronous group behaviors are also seen in yeast. For example, cells growing in a chemostatic, glucose-limited culture quickly synchronize their oxidative metabolism into regular, ∼3.5-hr cycles that are tied to the production of carbon monoxide from heme oxidation(Tu and McKnight, 2009, Tu et al., 2005). Acetaldehyde controls a quick, ∼1-minute-per-cycle oscillation of glycolytic intermediates in a population density-dependent fashion(Richard et al., 1996). At the whole-cell level, yeast grown on low-nitrogen medium develop pseudohyphae, or exhibit invasive growth with significantly elongated shapes(Gimeno et al., 1992). Such morphogenic adaptations are thought to help cells seek nutrients and spread outwards to explore fertile grounds. A network of signaling and cell adhesion pathways underlie this stress-stimulated differentiation(Kumar, 2021). Instead of an independent action by constituent cells within a colony, the differentiation from planktonic single cells to filamentous pseudohyphae utilizes a mechanism similar to quorum sensing(Albuquerque and Casadevall, 2012, Chen and Fink, 2006). Quorum sensing is a cell-to-cell communication system that coordinates individuals’ behavior based on the population density. In *S. cerevisiae*, 2-phenylethanol (PheOH) and tryptophol (TrpOH) have been identified as the major quorum sensing molecules(Chen and Fink, 2006, Albuquerque and Casadevall, 2012, Li et al., 2023) that stimulate pseudohyphal growth by triggering the expression of the flocculin protein Flo11p through Tpk1p of the cAMP-dependent protein kinase (PKA) signaling pathway(Chen and Fink, 2006). A third aromatic alcohol, tyrosol (TyrOH), appears to be less efficient for budding yeast, but facilitates dimorphic development of the pathogenic fungus *Candida albicans*(Chen et al., 2004). These alcohols are synthesized from the corresponding aromatic amino acids through the Ehrlich pathway during fermentation(Cordente et al., 2019), thereby tying metabolism to group behaviors. Besides pseudohyphal and invasive growth, diploid yeast cells can also go through meiosis to form spores, a dormant state that maintains viability through a variety of stresses for a long period of time(Huang and Hull, 2017). Apparently, yeast cells are equipped with a versatile repertoire of strategies to live through different environmental stresses.

In this study, we report a new developmental program of “viability resurgence in quiescent time” (VRQT) in *S. cerevisiae*. VRQT enables a small portion of stationary-phase cells to synchronously engage in and exit mitosis, resulting in a wave, sometimes waves, of population viability fluctuation. The synchronous mitosis reengagement in starvation is driven by the population density, a hallmark of quorum sensing. Indeed, quorum sensing aromatic alcohols enhance viability resurgence. Our findings provide new insights into the mechanisms of population rejuvenation and the extension of vitality under nutrient-scarce conditions.

## Results

Combination of starvation and heat stress expedites viability resurgence of aging yeast cells We previously reported that intracellular triacylglycerol (TAG) accumulation controls chronological lifespan in an energy expenditure-independent manner(Handee et al., 2016). TAG lipase knockout cells (i.e. *tgl3*Δ *tgl4*Δ) lack the ability to extract energy from the esterified fatty acids in TAG, therefore accumulating two-to three-fold excess TAG in the stationary phase. These “fat” cells exhibit significantly longer lifespan without a discernible defect in growth or survival. In contrast, cells lacking the TAG biosynthetic genes *DGA1* and *LRO1* cannot produce TAG and lose viability quickly after the culture enters the stationary phase, whereas the vegetative growth is largely unaffected. The stored TAG in cells is therefore regarded as a potent longevity factor, yet the molecular underpinnings remain to be delineated. In an attempt to identify genetic suppressors for the shortened lifespan of TAG deficient “lean” cells, we sought to define conditions that accelerated aging of yeast strains to expedite the subsequent screening and additional experiments. Among the different conditions tested, we found that growing cells at 37°C in synthetic complete (SC) medium resulted in shorter lifespan of all three strains while maintaining their differential viability in the stationary phase (Figure 1A). The time it took for each strain to drop their viability from ∼100% to 10% was 4, 5 and 8 days for the “lean”, normal, and “fat” strains, respectively. Under this expedited aging, cells manifested an elevation in viability after the overall survival fell to about 0.1% or lower – day 7, 11, and 20 for the three strains tested here. The net increase of clonogenesis (i.e., colony forming units, CFU) ranged from approximately 20-(wildtype) to more than 100-fold (fat strain) (Figure 1B). The resurgence in population viability of stressed yeast was also observed from cultures grown at the regular temperature of 30°C (Supplemental Figure 1). High-temperature growth thus accelerated the viability resurgence of starved cells but was not essential. Unless otherwise noted, we used 37°C culturing in SC medium as the standard protocol for faster turnaround of this yeast behavior. We refer to this phenomenon as viability resurgence (VR) and viability resurgence in quiescent time (VRQT) henceforth.

**Figure 1.**
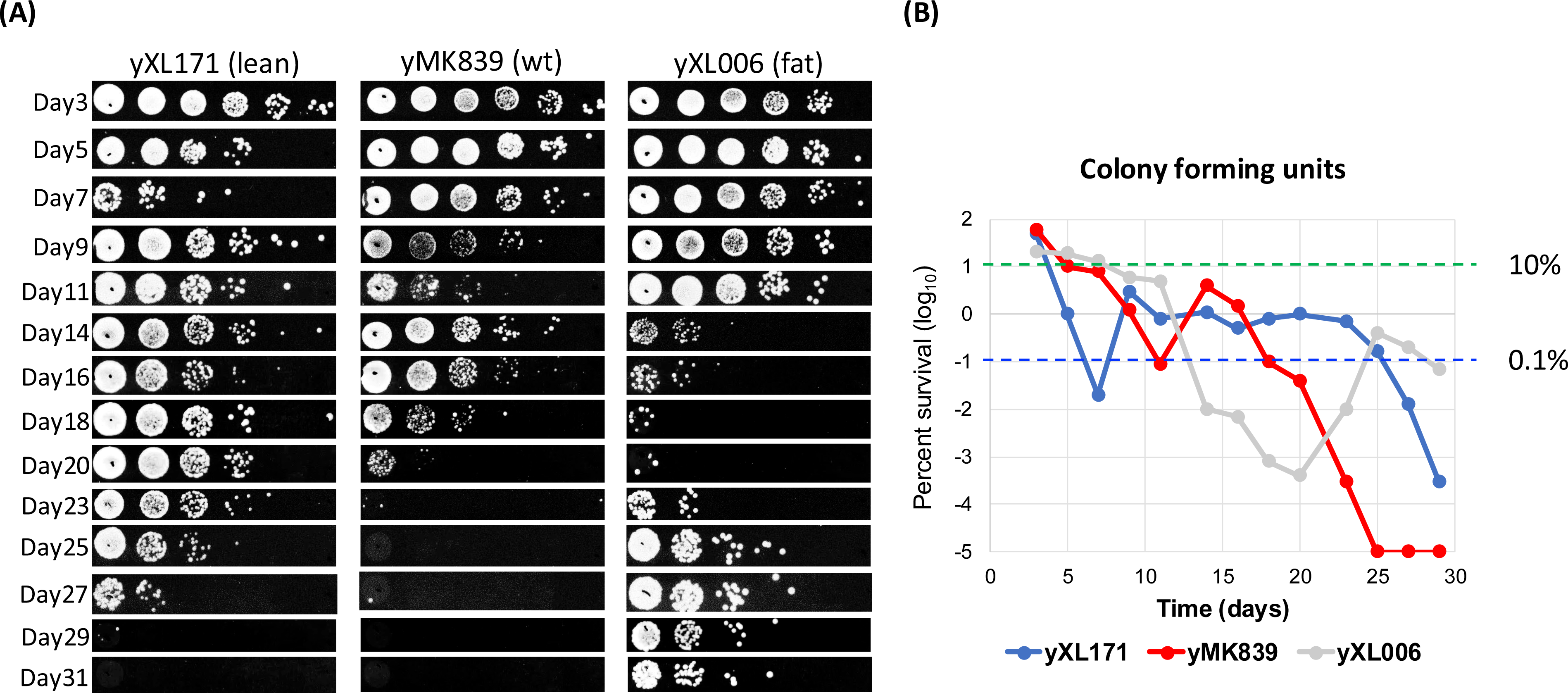
Combination of starvation and heat stress reveal viability resurgence of aged yeast cells. Yeast cultures from 37°C heat-stressed growth were collected and diluted in sterile water before they were spotted (left) or spread (right) onto fresh YPD plates to assess the population viability. Spot assays were done with 10-fold serially diluted cell suspensions. The first viability resurgence occurred at a time that conformed with the relative chronological lifespan of the three strains shown here. yXL171, yMK839, and yXL006 strains are triacylglycerol (TAG)-deficient, wildtype, and TAG-rich strains, respectively. The graph on the right depicts the colony forming units of the three strains from experiments done in parallel to the spot assays. Green and blue dashed lines represent 10% and 0.1% viability, respectively. Shown here are representative plots of minimally 3 independent tests of these strains.

We were curious as to how widely the VRQT phenomenon is distributed among *S. cerevisiae* strains. Additional laboratory strains (BY4742, W303, S288C, and Σ1278b), several prototroph wild strains from vineyard, woods or clinical samples, and a commercial wine yeast were tested for their survival curves in the stationary phase. Testing prototroph wild strains also helped rule out the influences on quiescence from auxotrophy and nutrient availability(Santos et al., 2021). All laboratory strains except W303 went through VR before dropping their overall colony forming unit to below 1 in a million (Supplemental Figure 2). Of the nine wild strains tested, the wine yeast, a clinical isolate (yKH073), and one from a vineyard (yKH077) displayed VR (Supplemental Figure 3). The remaining wild strains all showed very slow decline in their viability. At the end of the 36-day assay period, these strains exhibited an estimated CFU of 100 per million cells or higher (Supplemental Figure 3), consistent with our previous report that wild yeast strains have longer lifespan when tested at 30°C(Handee et al., 2016). Interestingly, the death curves of these wild strains were manifested by intermittent stabilization of viability (arrows, Supplemental Figure 3) that likely contributed to the extensive lifespan. We suspect that these were “mini-VR” that was sufficient to maintain the number of viable cells but not potent enough to cause a net increase of the overall viability (See Discussion). As to the commonly used laboratory strain W303, we noticed relatively moderate viability resurgence at 30°C (Supplemental Figure 4, days 44, 46, and 48), but this trait was curbed by incubation at 37°C. It is possible that this strain’s tolerance to sustained heat is impaired, resulting in the loss of viability before executing the VRQT program. While the detailed molecular and genetic reasons that underlie W303’s inablity to perform VRQT await further investigation, the frequent manifestation of VRQT among a wide spectrum of laboratory and wild yeast strains strongly suggests that VRQT is an innate survival program for yeast cells.

### Mutations do not likely cause viability resurgence during quiescent time

The VRQT behavior of yeast is reminiscent of the GASP phenomenon in prokaryotes, in which spontaneous mutations arise from cultures in prolonged stationary phase, resulting in steady viability for 16 days(Zambrano et al., 1993). Steinhaus and Birkeland(Steinhaus and Birkeland, 1939) first reported that *Sarcina lutea* and *Serratia marcescens* in laboratory batch cultures remained viable for two years. However, the yeast VRQT differs from these bacterial phenotypes in that VRQT involves fluctuation, not stabilization, of clonogenesis. Indeed, when colonies arose from VR (post-VR) were inoculated for a new round of growth, all 6 clones showed steady decline in survivorship after entering the stationary phase, but regained clonogenic capability after 5 to 9 days of growth (Figure 2A). The viability then decreased again, recapitulating the founding culture’s dynamic changes of survival rate. While some of these post-VR clones (e.g., No. 4 and 6) appeared to show slower decline of viability following VR (compare days 25 to 30), they still displayed a downward trend in survivorship. Critically, when the Area Under the Curve (AUC) of three pre- and six post-VR clones was calculated to compare the overall survival in 30 days, it is clear that the clones arisen from VR did not show discernible extension of the lifespan (Figure 2B; also see Supplemental Figure 5 for the three pre-VR samples for AUC calculation), arguing strongly against an advantageous mutation as the cause of VR.

**Figure 2.**
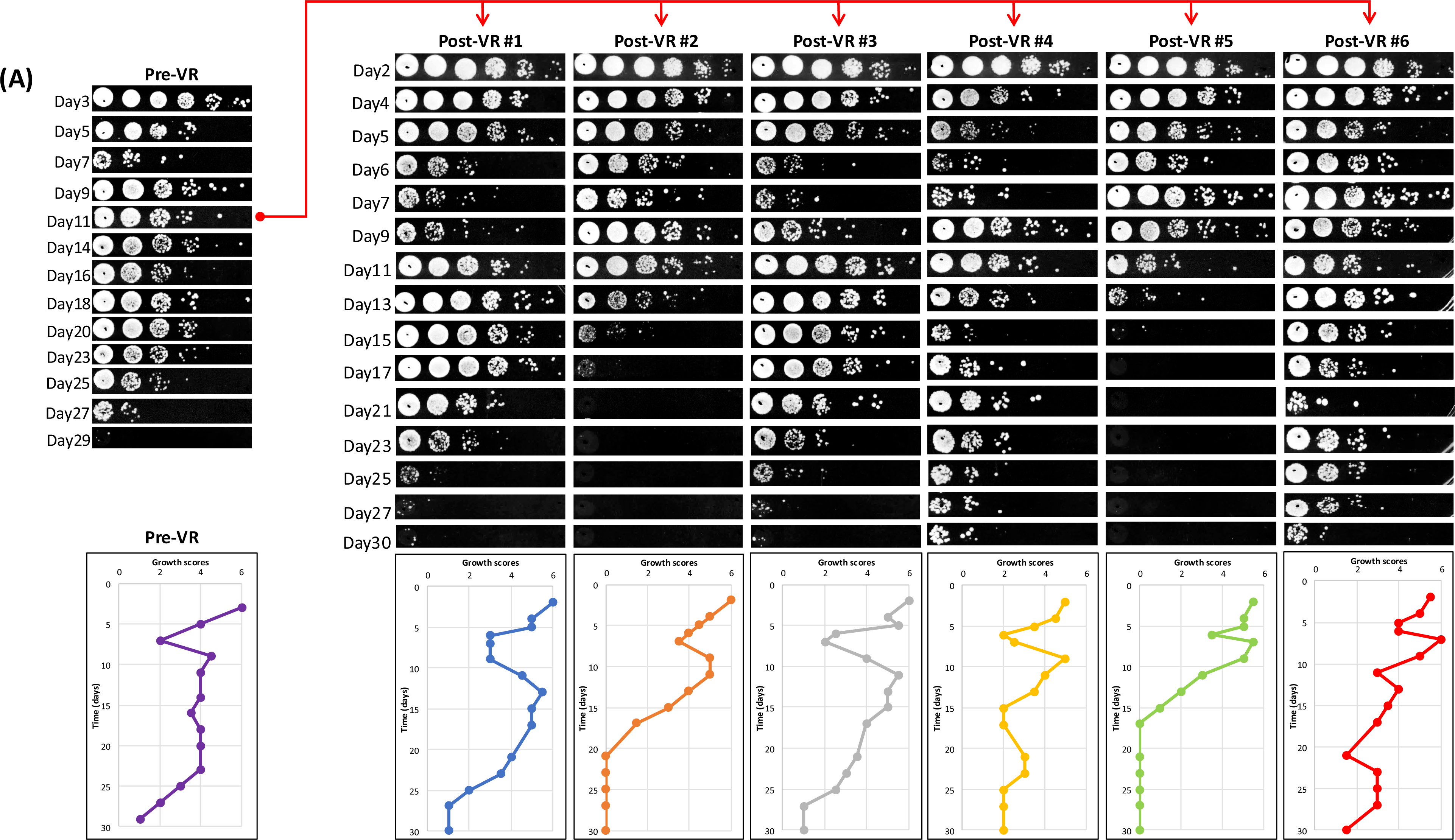
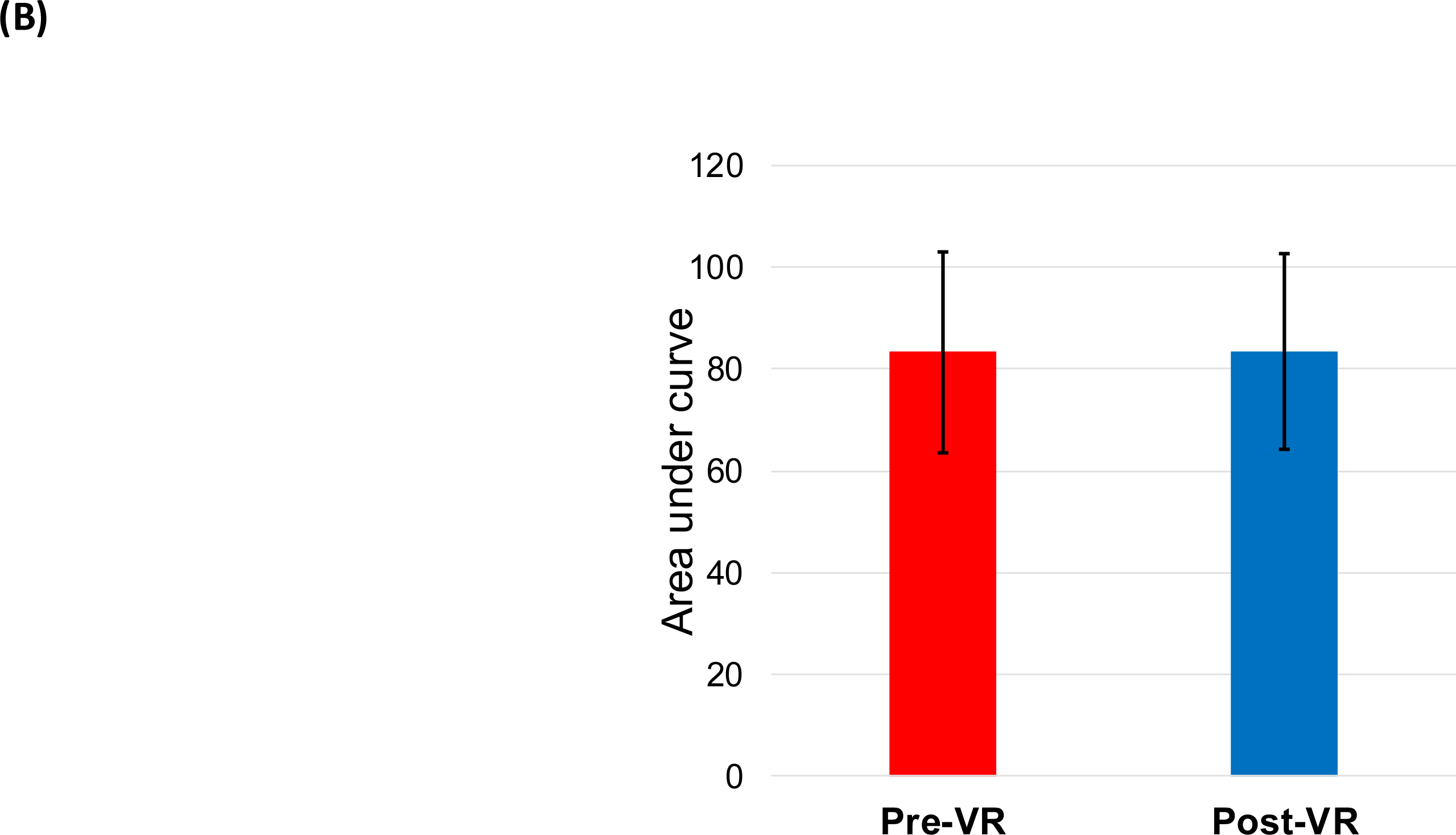
Cells re-grown from viability resurgence are not immortal, repeat the original fluctuation of survivorship in the stationary phase, and do not have a clear growth advantage over the founding cultures. (A) Post-viability resurgence (post-VR) single colonies from YPD plates were repopulated in SC medium for another round of chronological lifespan assessment under the heat stress. Shown are 6 post-VR clones (1 – 6) originally isolated from a day-11 yXL171 culture. The viability scoring charts are shown below the corresponding 30-day spot assay composites. See Materials and Methods for scoring rubric. All 6 post-VR clones showed similar rebound in viability. See Materials and Methods for scoring criteria. (B) Post-VR clones do not have a growth advantage over pre-VR cultures. The chronological lifespan of the pre- and post-VR clones were quantitatively compared by the Area Under the Curve (AUC) of each strain for 30 days. Shown are the AUC and standard deviations of three pre-VR biological replica (see Supplemental Figure 5 for the images), and six post-VR clones (as shown in Panel A).

### VRQT is mediated by an extracellular factor

Under certain conditions, yeast cells communicate with their neighbors. These include pheromones for mating(Herskowitz, 1995), aromatic alcohols for biofilm formation and pseudohyphal growth(Chen and Fink, 2006), purines for sporulation(Jakubowski and Goldman, 1988), ammonia for colony-to-colony communication(Palková et al., 1997), carbon monoxide to coordinate yeast metabolic cycles(Tu and McKnight, 2009), and acetaldehyde that synchronizes glycolytic oscillation(Richard et al., 1996, Weber et al., 2020, Aldridge and Pye, 1976). To assess whether VR might be mediated by an extracellular molecule, we conducted culture supernatant swap experiments. The spent medium of 7-day old lean (yXL171, early-VR) and fat (yXL006, late-VR) cell cultures was collected by centrifugation. Cell pellets were then washed and resuspended in the spent medium from the other strain or, as a control, in their own spent medium (Figure 3). Subsequent changes in viability were compared. In their own medium, the lean cells displayed the anticipated VR at day 9, whereas the fat strain viability declined slowly over the next 10 days. Strikingly, after re-suspending the fat cells in the day-7 lean cell spent medium, the recipient fat cells not only maintained a higher clonogenic capacity, they also exhibited VR after an additional 6 days (i.e., day 13 in Figure 3). Moreover, lean cells lost viability in the spent medium from the fat-cell culture. Importantly, the same lean-cell culture spent medium did not rescue the wildtype yMK839 cells at the same age (“yMK839/lean medium” in Figure 3). These results argued against the possibility that surplus nutrients released from the short-living lean cells supported the growth of cells indifferently. Instead, the results suggested that an extracellular signaling molecule(s) controlled the viability fluctuation of only those poised to respond to VR stimulation (detailed later). Indeed, if the medium swap was conducted four days later (D11 cultures), there was no exchange of the viability curves (Supplemental Figure 6), indicating that changes in the medium composition as well as the metabolic state of the cells are both important for successful execution of VRQT.

**Figure 3.**
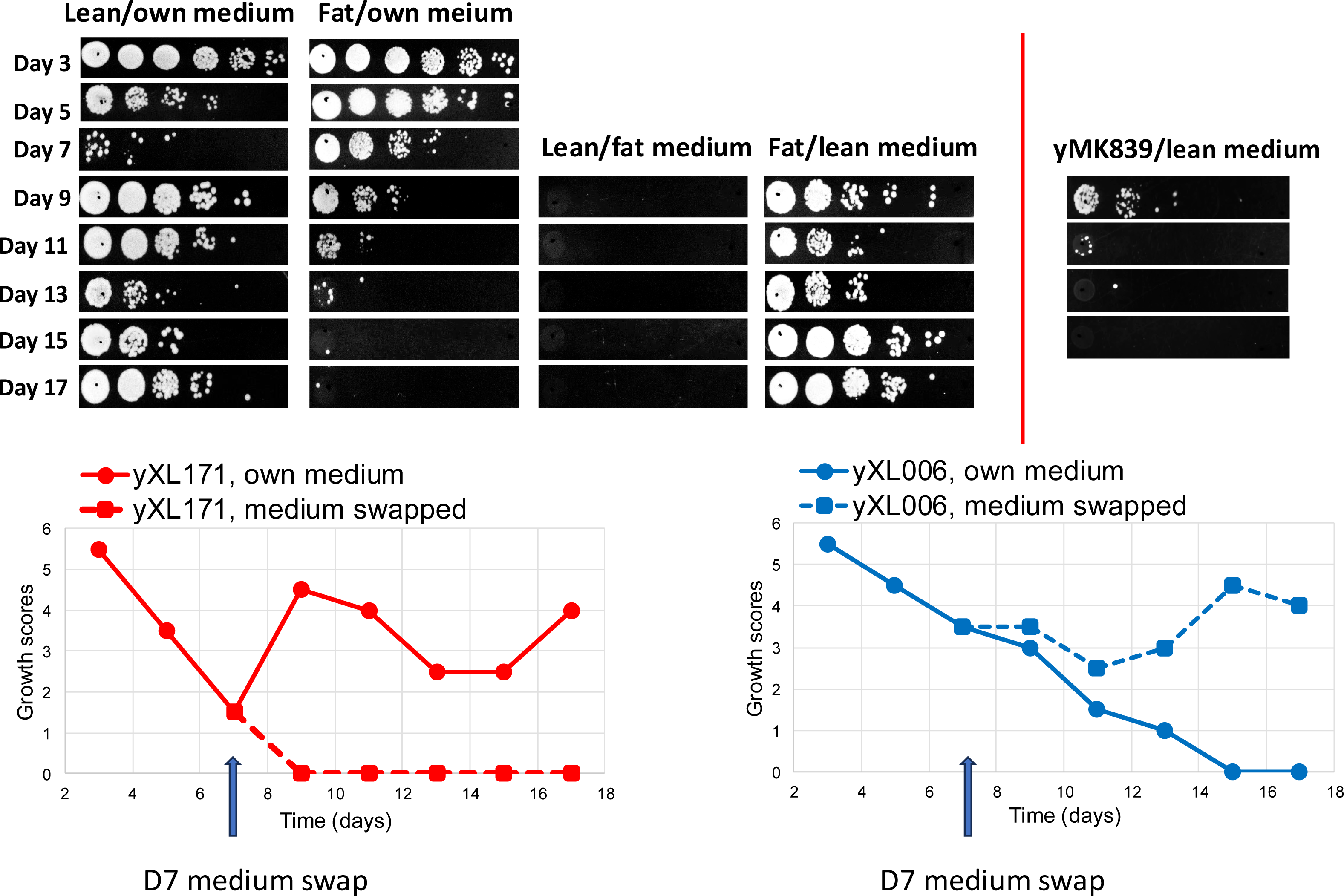
An extracellular factor mediates VRQT in yeast cultures. Spent medium from 7-day-old cultures of lean (yXL171) and fat (yXL006) cells were exchanged to test the effect on viability resurgence. Cells were separated from medium by centrifugation, washed with distilled water once, and resuspended in either their own spent medium or that of the other strain. Chronological lifespan was assessed by spot assays following medium exchange. Culture viability was measured every 2 days for 10 days.

To further characterize this inducing factor, we treated the spent medium from lean cells with DNase and RNase, or boiled the spent medium for 30 minutes to denature proteins, and found that none of these treatments impacted VRQT (Supplemental Figure 7). The VRQT inducer is therefore unlikely to be DNA, RNA, or proteins.

### VRQT is independent of known signaling pathways controlling cell growth and development

We next examined whether a specific signaling pathway for cellular growth or development was responsible for VRQT. Single-knockout strains of *tor1*Δ, *ras2*Δ, *kss1*Δ, *smk1*Δ, *far1*Δ, *slt2*Δ, *hog1*Δ, *atg8*Δ*, sod2*Δ, and *gpr1*Δ were constructed and tested for their survival in the stationary phase. All but two tested strains (*sod2*Δ and *gpr1*Δ) showed VRQT (Figure 4), suggesting that the mechanisms behind VRQT are separable from the conventional roles played by these genes in growth regulation and nutrient sensing (e.g., *TOR1*, *RAS2*)(Conrad et al., 2014, De Virgilio, 2012), development (e.g., *KSS1*, *SMK1*)(Ma et al., 1995, Krisak et al., 1994), stress response (e.g., *ATG8*, *HOG1*)(Tsukada and Ohsumi, 1993, de Nadal and Posas, 2022), cell cycle progression (e.g., *FAR1*)(Chang and Herskowitz, 1990), or cellular integrity (*SLT2*)(Martin-Yken et al., 2003). It is noteworthy that deleting the major caspase for apoptosis *YCA1*(Madeo et al., 2002) had no significant effect on VRQT, arguing against a close association between the VRQT survival program and apoptosis. The *ras2*Δ, *kss1*Δ, and *smk1*Δ cells reached the lowest level at day 17, 11, and 15, respectively, suggesting better maintenance of viability, i.e., lifespan extension, of these knockout strains when compared with the wildtype yMK839 tested at the same time. Conversely, deleting either *SOD2* (mitochondrial manganese-dependent superoxide dismutase)(van Loon et al., 1986) or *GPR1* (a plasma membrane G protein-coupled receptor)(Yun et al., 1997) diminished VRQT greatly. Table 1 shows that the average frequency of VR of the tested strains was as high as 79%, but the *sod2*Δ and *gpr1*Δ strains showed VRQT in 0 out of 3 trials, and 4 out of 18 trials, respectively. We noticed that the fat strain (*tgl3*Δ *tgl4*Δ) and the *atg8*Δ mutant exhibited VR at 50% of the tests. Whether increased TAG storage and reduced autophagy inhibit VRQT awaits further investigation.

**Figure 4.**
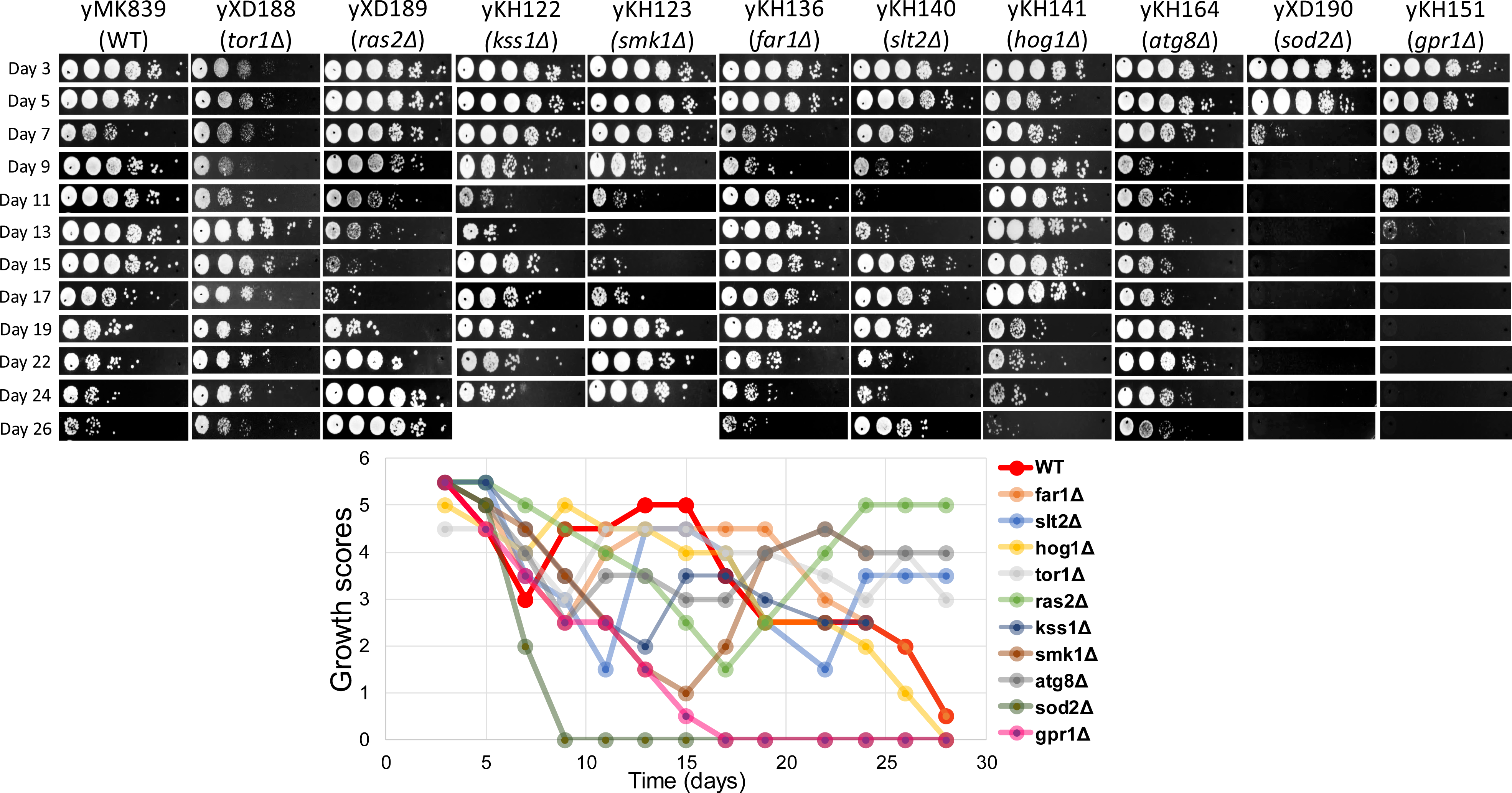
VRQT is independent of several signaling pathways key to cellular growth and development. Single-knockout strains as indicated above each column of images were constructed from yMK839, a standard "wildtype" strain in this study to assess the genetic interaction between VRQT and some of the well-studied pathways for yeast growth and development, as well as several that are known for their contribution to chronological lifespan control. Additional knockouts were made in the yXL171 background. Of all the knockout strains tested, only *SOD2* and *GPR1* deletion strains show severely impaired viability resurgence. See Table 1 for summary.

**Table 1.**
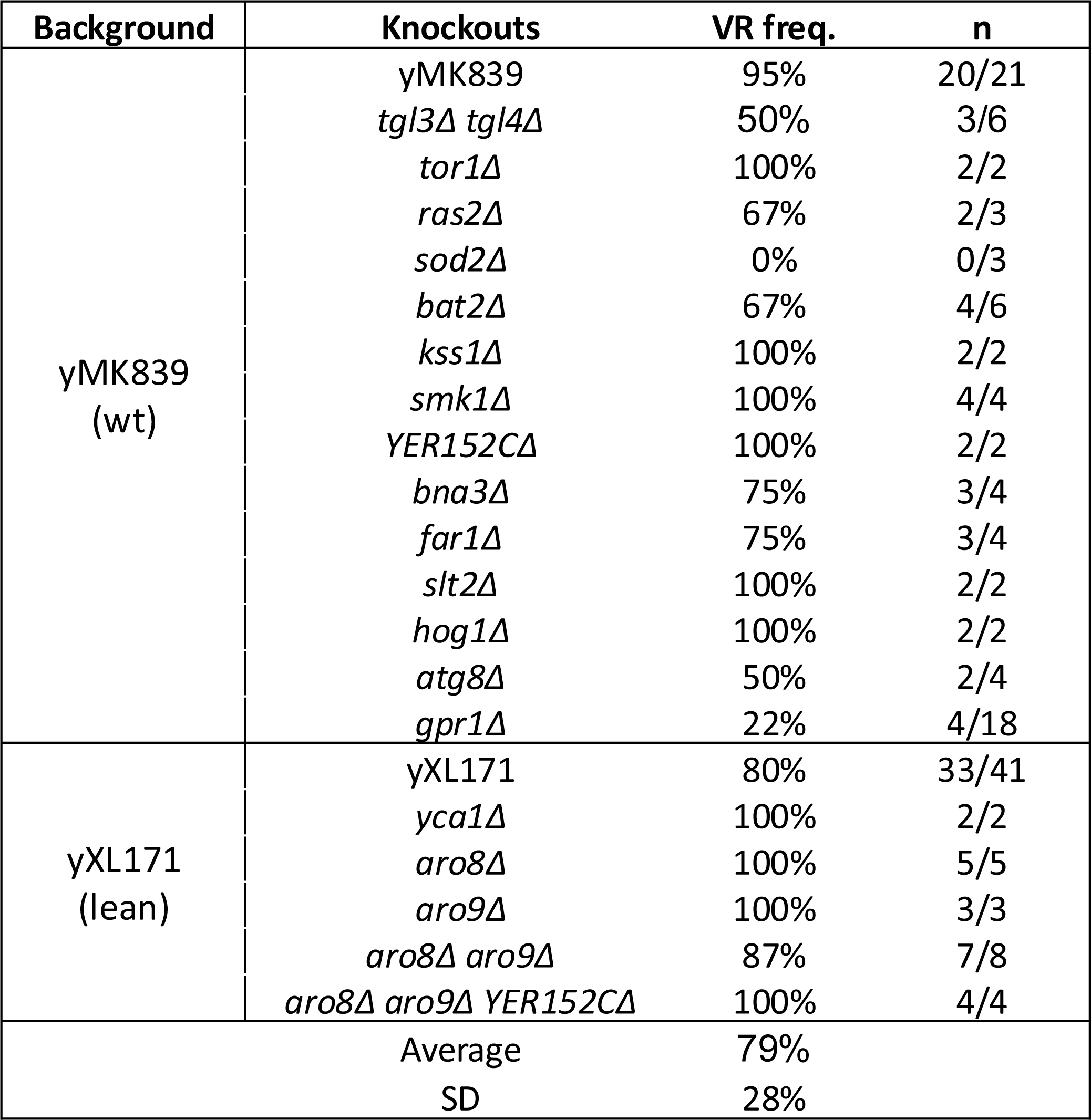
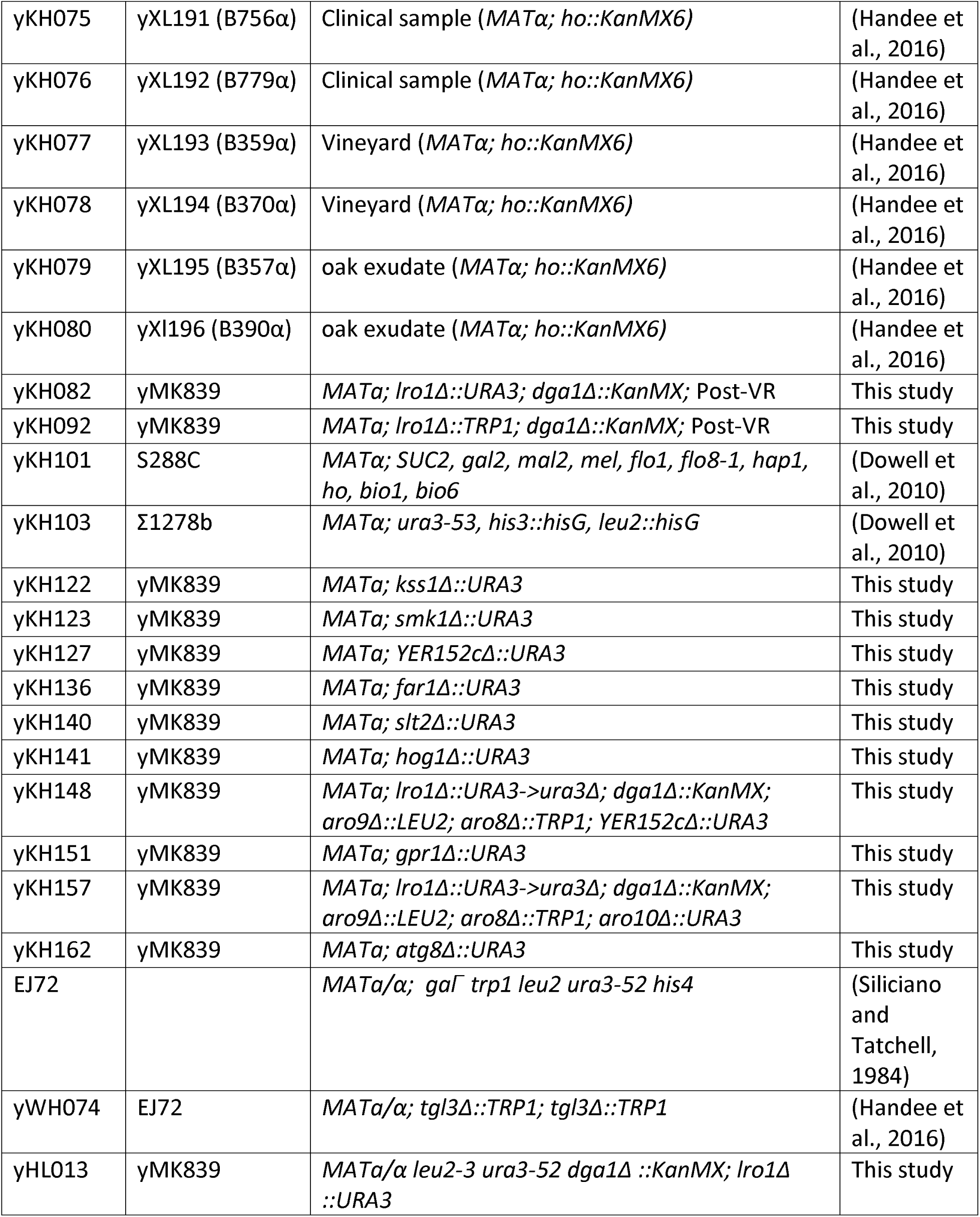
Yeast strains. The viability resurgence in quiescent time requires *SOD2* and *GPR1* genes, but is independent of many pathways that control cellular development and growth.

Close inspection of the spot assays revealed that, in most of the strains, the decline of the population viability followed a relatively shallow slope. However, *sod2*Δ cells consistently showed abrupt collapse between day 5 and day 7 (see, yXD190 in Figure 4), suggesting that the lack of the mitochondrial superoxide dismutase Sod2p causes sudden death so cells may not have sufficient time to instigate their VRQT program. On the other hand, the *gpr1*Δ cells (yKH151) showed similar kinetics of viability loss as most of the other strains, but a low frequency of VR (4 VR out of 18 trials). Gpr1p is a non-essential G protein-coupled receptor (GPCR) that constitutes one of the two branches of the cAMP/PKA pathway for glucose sensing(Busti et al., 2010) for cellular growth and division. Whether this GPCR is responsible for sensing and conveying the VR inducing signal is an intriguing question to be examined.

Besides genetic knockouts in a haploid background, we examined whether conditions associated with VRQT activated sporulation in diploid cells. The homozygous diploids of normal, lean and fat strains were first confirmed for their ability to exhibit VRQT (Supplemental Figure 8A). At day 23, each of these three strains was sampled and stained by DAPI, and the number of nucleus of each cells was visualized by epifluorescence microscopy. The diploid fat cells (yWH74) were also incubated in a regular sporulation medium (SPM) for 7 days before DAPI staining. While cells sporulated and formed tetrads efficiently in SPM (red arrows, Supplemental Figure 8B), none of the three diploid strains showed tetrads when collected from the stationary-phase SC medium cultures. Together with the genetic knockout data from Figure 4, we conclude that VRQT represents a stress-responding program that is carried out by a mechanism distinct from many well-characterized pathways controlling yeast development and stress responses.

### VRQT depends on the population density and involves nearly synchronous cell division

The increased clonogenesis during VRQT may arise from cell division in spent medium, or from conversion of senescent cells back to quiescent cells(Li et al., 2018). In theory, the latter can be tested by separating quiescent (Q) from non-quiescent (NQ) cells with isopycnic centrifugation in a Percoll gradient(Allen et al., 2006). However, we found that late stationary-phase yeast cells lost the Q/NQ density differentiation resolvable by Percoll or iodixanol(Allen et al., 2006, Quasem et al., 2017). Instead, we examined the first hypothesis, that is, whether propagation of cells in a spent medium was an underlying cause for VRQT. To capture the very small dividing population, if existent, we prepared 2% agarose slabs made with the spent medium from a 9-day old lean cell culture (see Figure 5A for the experimental design). Cells of different ages were centrifugation-collected, washed, and suspended (3-fold serially diluted) in water before spotting on the spent medium/agarose slab (SMAS). These inocula were put back to 37°C incubation. Colony formation on this agarose slab would provide definitive proof for cell division in spent medium. After 2 to 3 days of 37°C incubation, yeast inocula, along with the agarose slab underneath, were transplanted to a fresh YPD plate of 30°C growth to mimic the standard YPD spotting assays that quantified clonogenesis (e.g., Figure 1). Nutrients in YPD diffused upwards into the agarose slab and triggered mitosis-capable cells (i.e., quiescent cells) to grow, consequently revealing the overall viability.

**Figure 5.**
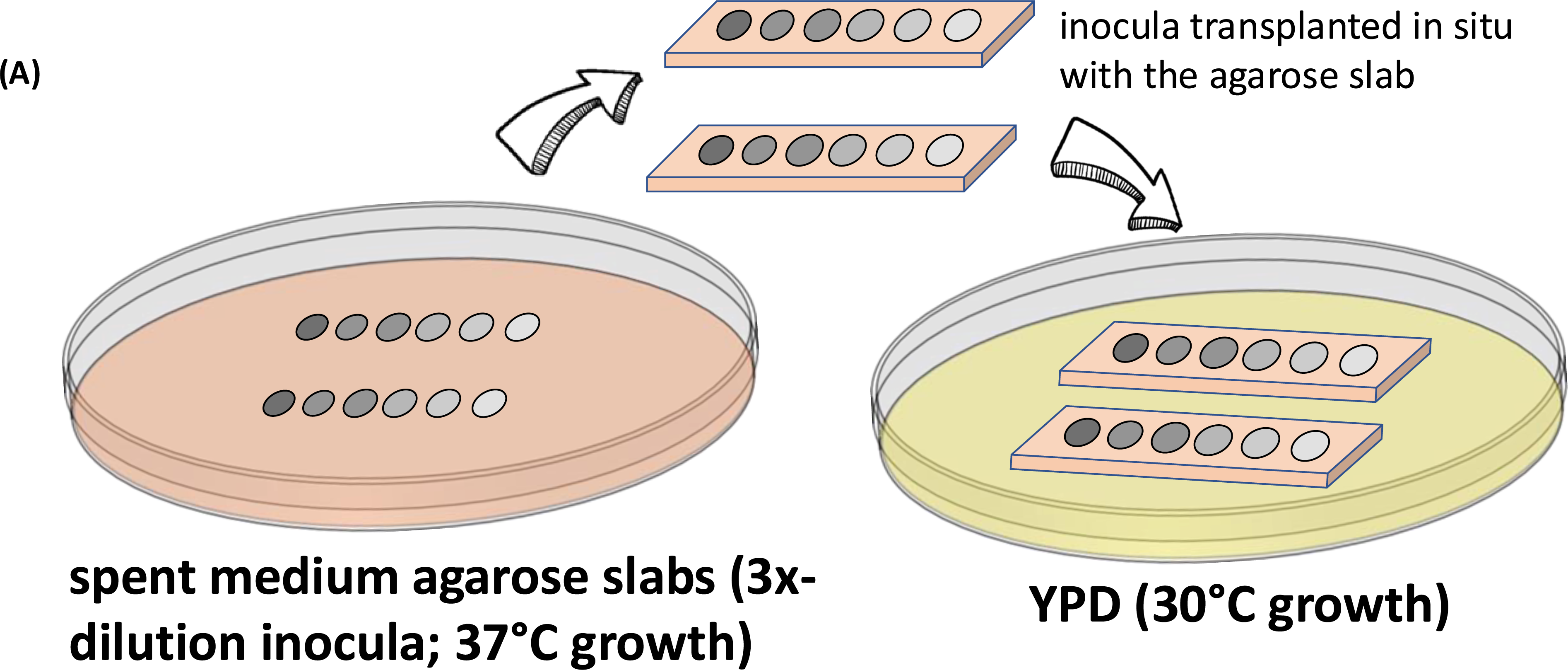
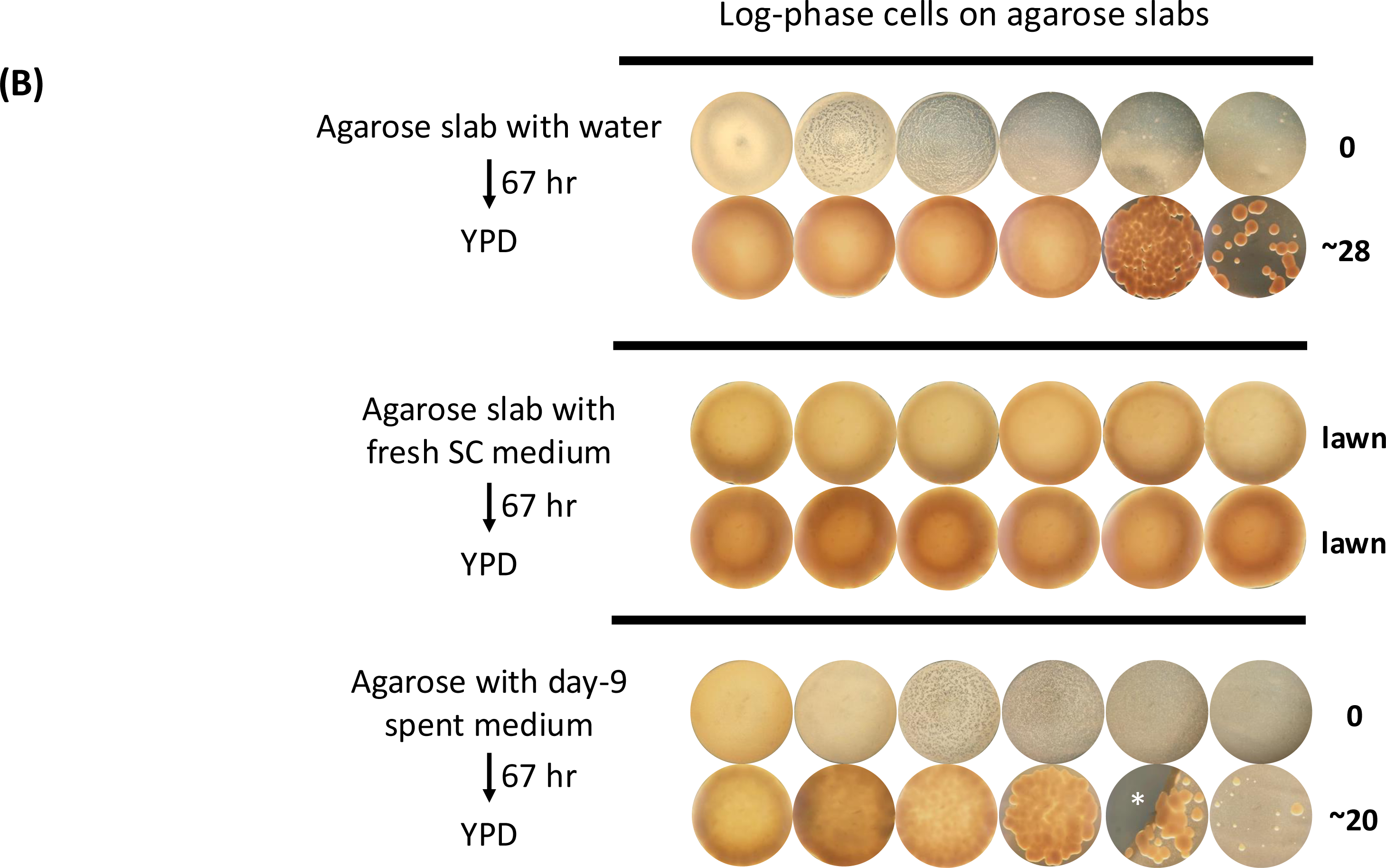
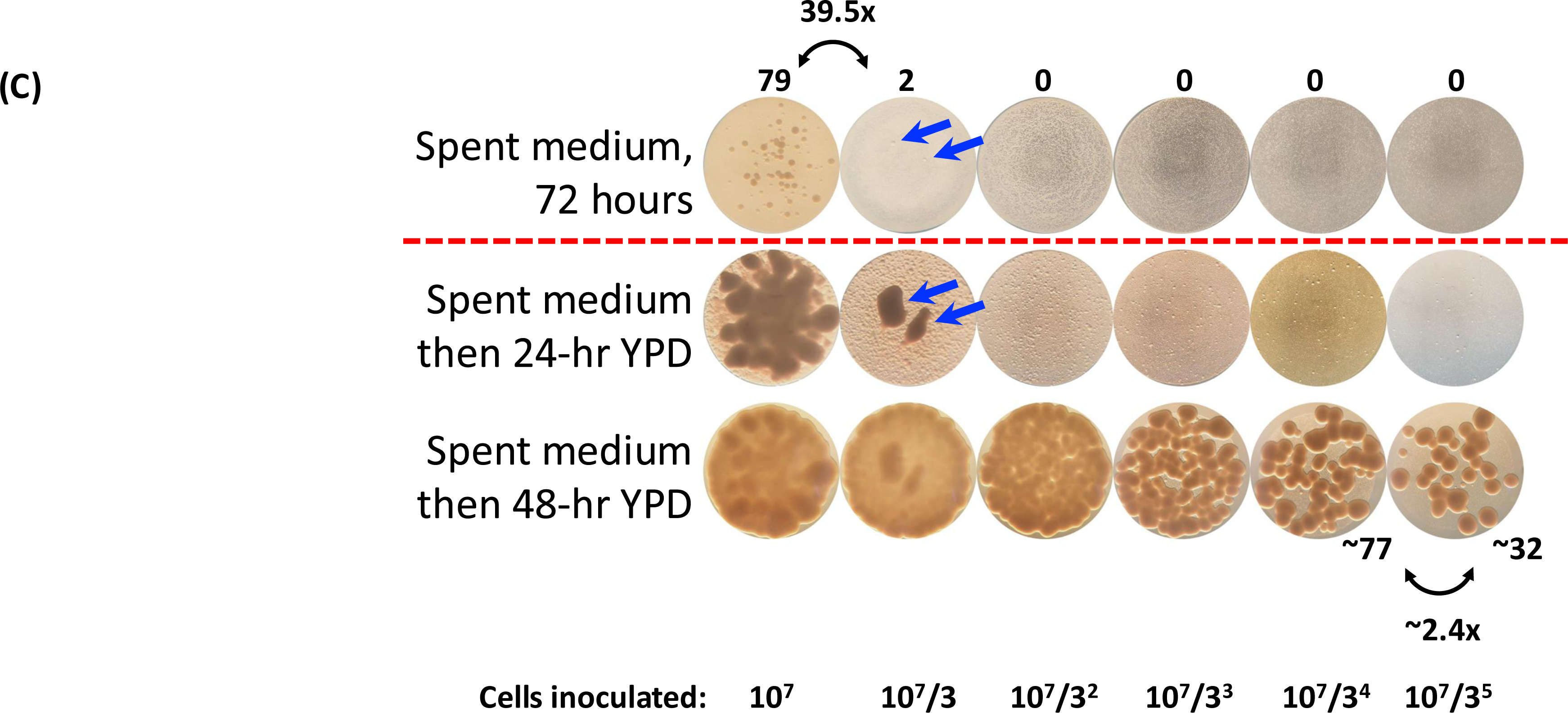
VRQT is a result of nearly synchronous cell division linked to the population density. (A) Schematic of <u>s</u>pent <u>m</u>edium <u>a</u>garose <u>s</u>lab (SMAS) assay experimental design. Cells were 3-fold serially diluted in sterile water and spotted (5 µL each) onto 2% agarose slabs that were prepared from spent medium. Inocula on the agarose slabs were incubated at 37°C to mimic the growth conditions for VRQT. After 2 to 3 days, the cell inocula were photographed before the agarose slabs with the inocula were excised and transplanted to fresh YPD plates for growth at the normal 30°C incubation for 2 days to reveal viable and mitotically capable cells. The second phase of the SMAS assay mirrors the traditional YPD spot assays. (B) Feasibility of the SMAS assay using log-phase cells and agarose slabs made of different media: distilled water, fresh SC medium, or day 9 yXL171 lean cell spent medium. 5 µL of 3x serially diluted yXL171 log-phase cells were spotted to these slabs and incubated at 37°C for 3 days, followed by transplantation of slabs to YPD for two more days at 30°C. Each inoculum was photographed under a dissection microscope. The numbers of discernible colonies on the highest dilution are shown on the right of the inoculum image. Asterisk in the image of the bottom row, second from the right, indicates a broken slab during the experiment. (C) 12-day-old yXL171 cells were spotted onto agarose slabs prepared from day 9 spent medium from a separate yXL171 culture. 3x serial dilution started with ∼10,000,000 cells on the leftmost inoculum. The agarose slabs were incubated at 37°C for 72 hours, then transplanted to grow on fresh YPD plates for 48 hours. Blue arrows point to the two “primary” colonies on the spent medium agarose slab, which continued to grow on YPD plates. Many “secondary” colonies only appeared after transplantation to YPD. Colony numbers of selective inocula are shown by the photo. Note the strong disparity between the number of the primary colonies and the 3x inoculum dilution. The number of the secondary, YPD only, colonies was closer to the 3x dilution of the inocula.

To ascertain the feasibility of the SMAS assay, we first used log-phase cells (Figure 5B). An agarose slab made of water was prepared which did not have nutrients needed to support cell division. After transplantation to YPD, colonies were abundant, verifying the upward diffusion of nutrients from YPD to the slab. As expected, robust growth of log-phase cells was seen on agarose slabs made of SC medium even before the transplantation to YPD (middle row, Figure 5B). Spent medium (bottom row), just like water, could not support clonogenesis of log-phase cells before exposure to YPD. These spent-medium-then-YPD colonies were significantly smaller than those grown from the water/agarose slab-then-YPD counterparts, suggesting that logarithmically growing cells, unlike quiescent cells that are resistant to many stresses, were hindered by the 9-day culture spent medium containing harmful metabolites such as acetic acid and reactive oxygen species(Burtner et al., 2009).

In contrast to the log-phase cells that failed to form colonies on spent medium before transplantation to YPD, cells from a 12-day old stationary-phase culture grew colonies indicative of proliferation on the spent medium (arrows, Figure 5C). After transplantation to YPD and 24-hr incubation, the small colonies grew proficiently while additional small colonies appeared. This two-stage growth test provided solid evidence for the existence of two populations of viable cells in an aged culture: the first one consists of cells that divide under the same deprived and stressful condition. The second population consists of quiescent cells that stay dormant until exposure to fresh medium and then begin to divide. Further inspection of the numbers of the colonies before YPD exposure revealed an important population behavior consistent with the phenotype of quorum sensing, that is, a population member density-dependent collective action: Whereas cells from the 12-day old stationary-phase culture were inoculated for the SMAS assay at three-fold serial dilutions, there was a 79-to-2, 40-fold difference in the colony numbers for the two densest inocula. No colony was seen in subsequent, higher dilutions. In contrast, the number of colonies that emerged after transplantation to YPD decreased as expected for the 3-fold serial dilution through the entire inoculation regime. For example, there were approximately 77 and 32 colonies on YPD for the two highest dilutions, much closer to the anticipated ratio of 3-to-1. The population-dependent colony formation on SMAS was a highly reproducible behavior. Supplemental Figures 9 and 11 show additional, independent examples of this phenomenon.

To determine how many generations VR cells went through to generate discernible colonies on spent medium, we plucked 8 colonies from 4 separate inocula from a SMAS and suspended the cells for cell counting. We picked colonies with a diameter of about 500 µm (451 – 711 µm, mean = 527 µm), and the average number of cells from each colony was found to be 43,516, which corresponded to about 15 divisions (Supplemental Figure 10). Typically, each boom of clonogenesis on SMAS did not last any more than two days (our standard increment for viability and SMAS assessments) (also see Figure 7 and Supplemental Figure 12 later), which indicated a 3.2-hr period for each division, assuming 100% of cells in the colony are dividing non-stop in the spent medium. Yeast cells in a colony exhibit different physiology, depending on their location in the colony(Cap et al., 2012) and the colony’s age(Meunier and Choder, 1999). The 3.2-hr growth rate on SMAS therefore is considered reasonably close to the maximal division rate of 90 minutes in a rich medium(Meunier and Choder, 1999). We therefore infer that VRQT involves largely synchronous cell division of the capable cells.

To further correlate SMAS clonogenesis to viability resurgence as revealed by YPD spot assays, we conducted a detailed comparison of four independent samples of yMK839 (normal TAG, 1 culture) and yXL171 (TAG-deficient lean strain, 3 cultures). Samples were collected every 24 hours and subjected to SMAS and YPD spot assays concurrently. Scoring results clearly demonstrated that each viability resurgence or stabilization was accompanied by colony formation on SMAS (blue and orange shades, respectively; Supplemental Figure 11). Thus, it is concluded that concerted cell division, which can be captured by the SMAS assay, is the underlying cause of VRQT.

### Quorum sensing molecules modulate kinetics of VRQT

The population density-dependent cell division on spent medium conforms to the canonical behavior of quorum sensing. Quorum sensing is a means of intercellular communication widely employed by microorganisms to perform collective actions such as emission of bioluminescence, swarming migration, biofilm formation, and virulence factor production(Bassler, 2002, Yashiroda and Yoshida, 2019). In budding yeast, the aromatic alcohols 2-phenylethanol (PheOH) and tryptophol (TrpOH) are the quorum sensing molecules (QSMs) that trigger pseudohyphal growth of certain diploid strains starved for nitrogen on a solid agar plate(Chen and Fink, 2006). The third aromatic alcohol, tyrosol (TyrOH), does not appear to have this effect under the same condition(Chen and Fink, 2006), but can stimulate dimorphic development in *Candida albicans*(Chen et al., 2004). Because of the apparent population density-dependent division of aged cells on spent medium, we tested whether supplementing the spent medium with aromatic alcohols could influence VRQT. SMAS colonies from the inocula of a 12-day-old culture were examined after 43 hours of incubation at 37°C, followed by transplantation to YPD plate for an additional 18 hr (Figure 6A). Without aromatic alcohol supplementation, colonies were not observed on the spent medium in this experiment (top row; number in circle represents the corresponding colony number before transplantation). At 100 µM, all three aromatic alcohols stimulated yeast cells to divide and form microcolonies.

**Figure 6.**
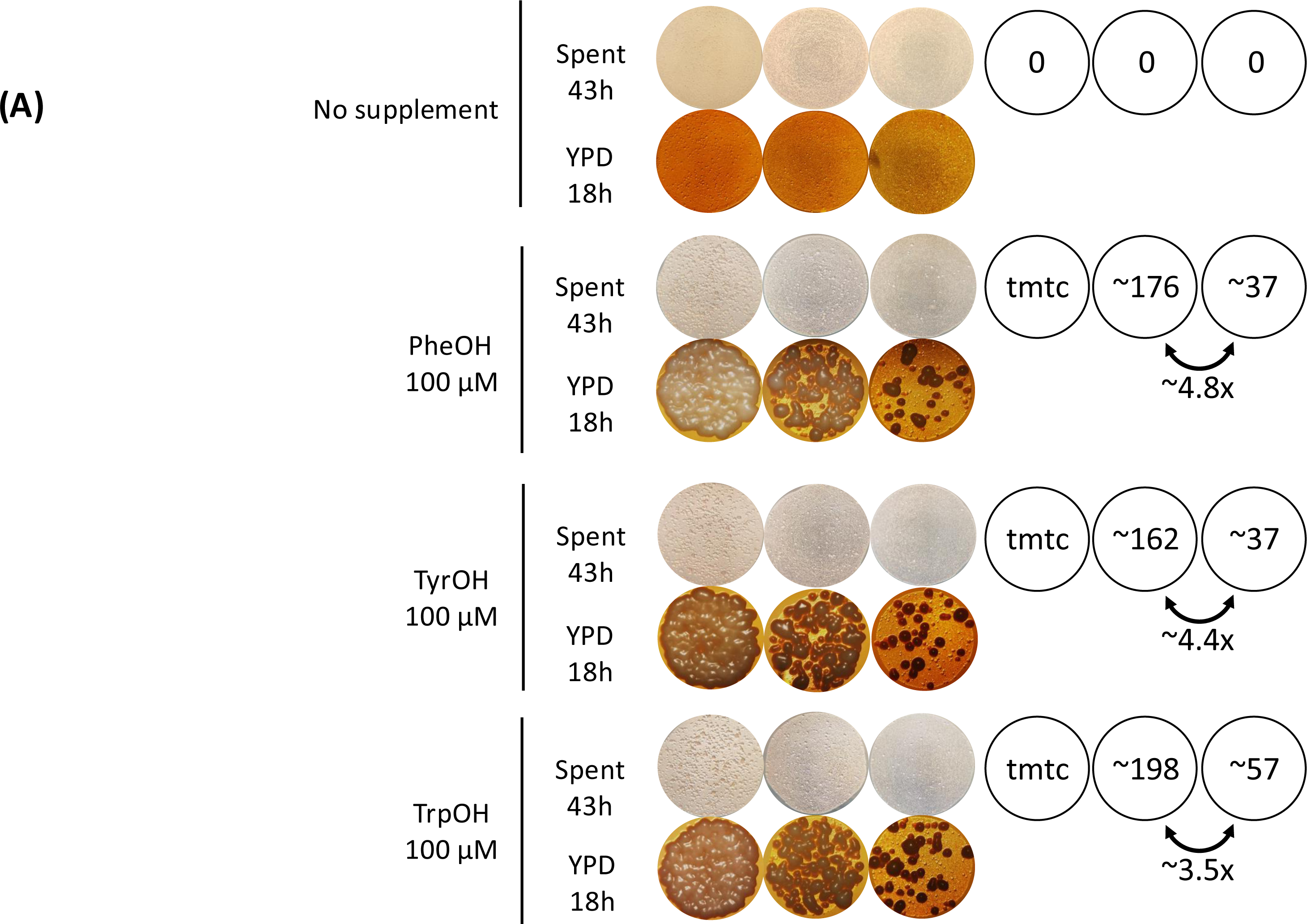
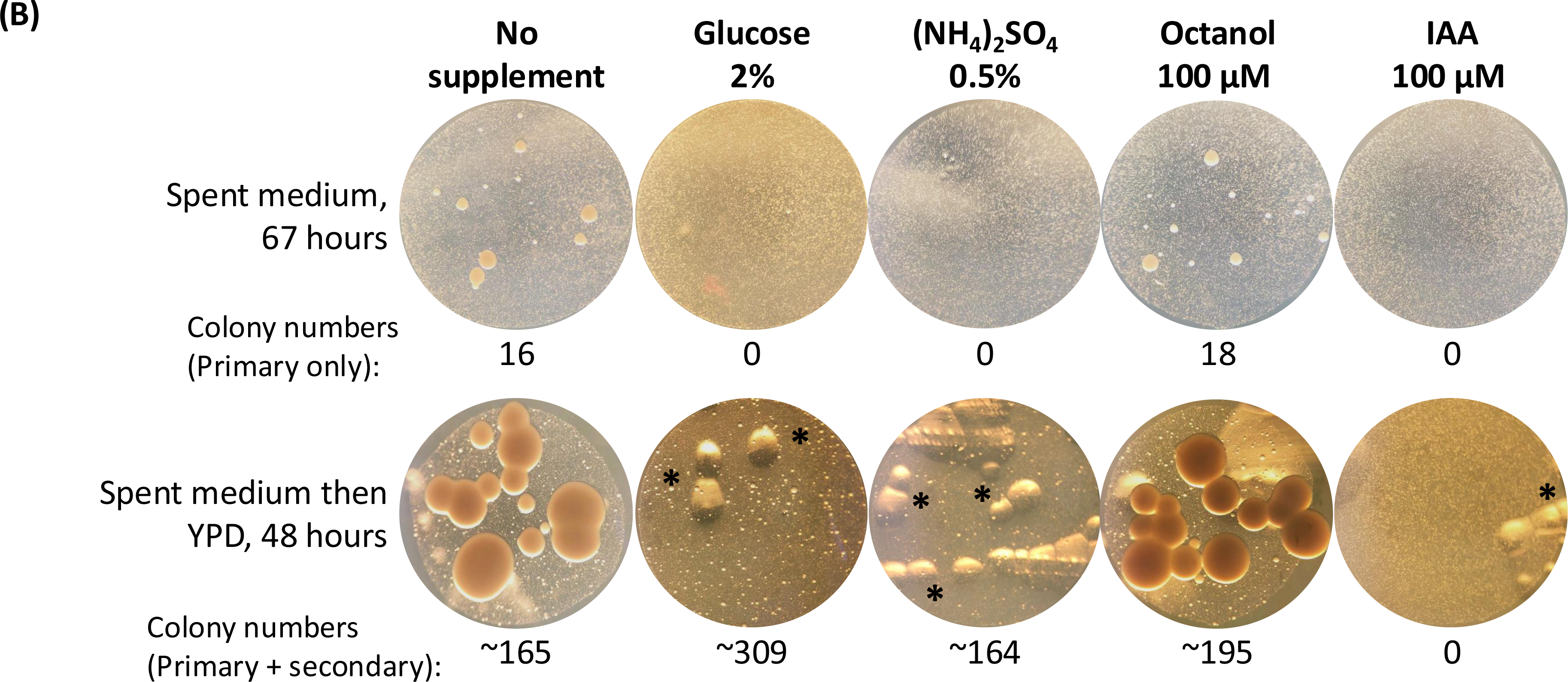
Quorum sensing aromatic alcohols stimulate cell division that underlies VRQT. (A) Supplementing aged cells with aromatic alcohols stimulates colony formation in a pattern that is less sensitive to the population density. 3x serially diluted 12-day-old yXL171 cells were spotted onto SMAS of yXL171 day 9 spent medium. Each agarose slab has been supplemented with 100 μM of 2-phenylethanol (PheOH), tyrosol (TyrOH), or tryptophol (TrpOH) before cell suspensions were inoculated. The yeast inocula were incubated at 37°C for 43 hours, then slabs were transplanted onto fresh YPD plates for another 18-hr incubation at 30° to better visualize the colonies that had formed on SMAS but with minimal secondary colonies forming at this time point (see Figure 5C). Colony numbers are shown on the right; tmtc = too many to count. (B) VRQT stimulation is specific to aromatic alcohols. Supplementing with glucose, ammonium sulfate, octanol, or indole acetic acid (IAA) does not stimulate VRQT. Experiments in panel A were done with a supplement of 2% glucose, 0.5% ammonium sulfate, 100-µM octanol, or 100-µM IAA (indole acetic acid). None of these four chemicals stimulated colony formation. Air bubbles (asterisks) were inadvertently introduced to the YPD transplantation plates.

**Figure 7.**
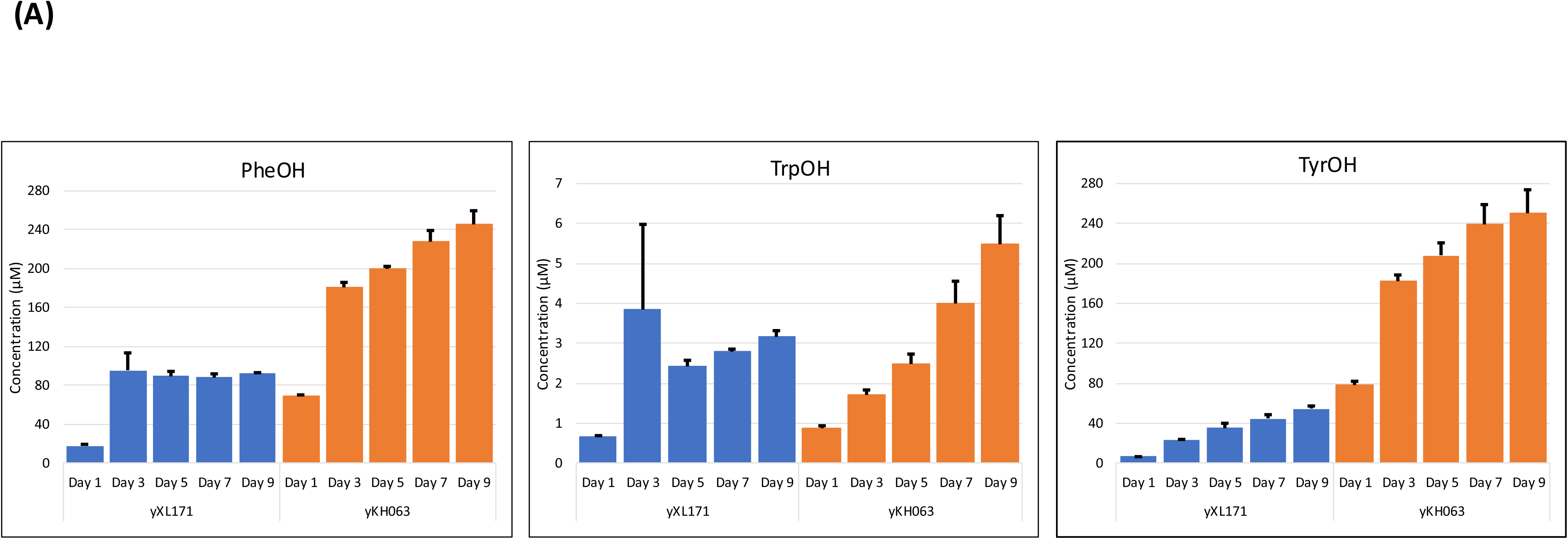
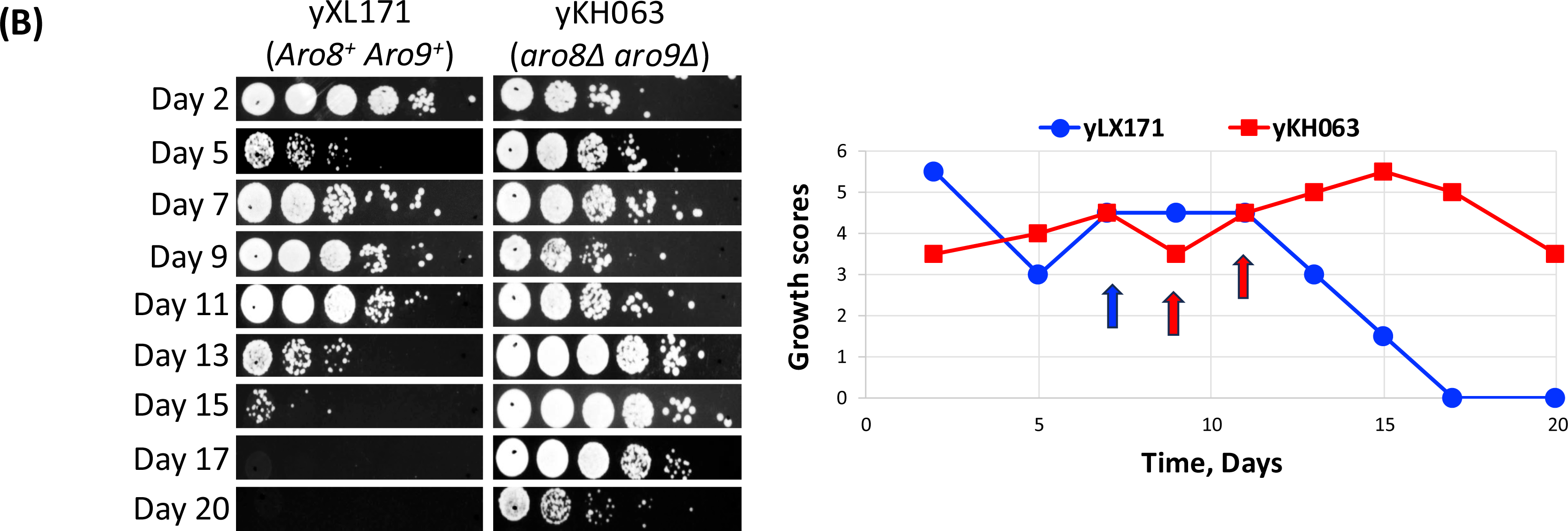
Deleting *ARO8* and *ARO9* enhances aromatic alcohol production and cell survivorship. (A) HPLC quantification of the aromatic alcohol concentrations in the medium of the parental (blue bars) and *aro8*Δ *aro9*Δ (orange bars) strains grown for 9 days. Abundance of all three aromatic alcohols increased when cultures grew older. Deleting both *ARO8* and *ARO9* further elevated these alcohols. Note that TrpOH quantification is presented at a different scale at y axis. (B) Viability curves of *Aro8*^+^ *Aro9*^+^ and *aro8*Δ *aro9*Δ cells. Note the apparent growth advantage of the *aro8*Δ *aro9*Δ cells (red curve). Blue and red arrows indicate the time when these two strains showed colonies on spent medium agarose slabs (See Supplemental Figure 14 for the complete set of images).

At 10 µM, TrpOH stimulated cell division, but the other two did not (Supplemental Figure 12). The critical concentration to stimulate VR, therefore, was no more than 10 µM for tryptophol and between 10 and 100 µM for 2-phenylethanol and tyrosol. The strong VR inducing activity of TrpOH is consistent with the HPLC quantification results (Figure 7A, blue bars) in that this alcohol was the least abundant among the three aromatic alcohols in the spent medium. Moreover, it is also worth noting that aromatic alcohol supplementation diminished the population-density dependence for colony formation on the spent medium (compare Figures 6A and 5C), suggesting that aromatic alcohols were key to the quorum sensing-like VR of aged yeast cultures.

Aromatic alcohols likely acted as quorum sensing signaling molecules rather than supplying nutrients that supported cell division. In a separate supplementation experiment in which stationary-phase cells were spotted to SMAS supplemented with 2% glucose, 0.5% ammonium sulfate, or 100 µM 1-octanol, the 2-phenylethanol equivalent without the benzene ring, formation of microcolonies was not stimulated by any of these treatments (Figure 6B). In fact, glucose and ammonium sulfate, the major carbon and nitrogen nutrients in SC medium, prevented cell division on the spent medium. Interestingly, the number of secondary colonies that formed after transplantation to YPD nearly doubled with 2% glucose supplementation; ammonium sulfate did not have this effect. These observations suggested that the overall viability might be extended by additional carbon source, but the ability to execute mitosis was not stimulated, rather it was suppressed by these two nutrients. We also examined the outcomes of supplementing with indole-3-acetic acid (IAA), the fully oxidized product from tryptophan(Cordente et al., 2019). Contrary to TrpOH, IAA not only inhibited microcolony formation but also reduced the overall viability to less than 1 in 10 million on YPD (see Supplemental Figure 13 for a complete set of images). IAA is synthesized by yeast in tryptophan-dependent and -independent pathways(Rao et al., 2010). Both TrpOH and IAA stimulate invasive growth(Prusty et al., 2004). The observations that tryptophol induced whereas IAA inhibited stationary-phase cell division demonstrate the distinctions between VRQT and pseudohyphal growth.

### Deleting *ARO8* and *ARO9* elevates aromatic alcohols, enhances VRQT, and affords growth advantage in the stationary phase

Aromatic alcohols are catabolic products of the corresponding aromatic amino acids.

Under the typical laboratory growth conditions, conversion of these amino acids into alcohols is through the Ehrlich pathway that synthesizes fusel aldehydes(Cordente et al., 2019)(See Supplemental Figure 15 for the abbreviated pathway). Fusel aldehydes can be reduced to form fusel alcohols (e.g., the three aromatic alcohols described in this work) or oxidized to generate carboxylic acids such as IAA. If aromatic alcohols are key to VRQT, manipulating their biosynthetic enzymes would alter the abundance of such metabolites and, correspondingly, the prevalence of VRQT. To test this notion, we knocked out *ARO8* and *ARO9*, which encode the two reversible aromatic aminotransferases of the Ehrlich pathway(Iraqui et al., 1998). While Chen and Fink(Chen and Fink, 2006) showed that deleting these two genes reduced the conversion of radiolabeled aromatic amino acids to the corresponding alcohols by actively growing cells, Zhang et al showed that the steady-state levels of such alcohols had increased in the knock-out strains during biofilm formation(Zhang et al., 2021), highlighting the reversible nature of these two aminotransferases. Given the seemingly opposite outcomes of deleting these two genes, we reasoned that up- or down-regulation of the abundance of aromatic alcohols resulting from deleting *ARO8* and *ARO9* would provide us a means to assess the contribution made by these alcohols to VRQT. Indeed, HPLC quantification of the aromatic alcohols secreted by the parental yXL171 and the *aro8*Δ *aro9*Δ yKH063 strains showed that the latter strain produced higher steady-state levels of all three aromatic alcohols in the stationary phase (Figure 7A), consistent with the model presented by Zhang et al (see Discussion for details)(Zhang et al., 2021). Importantly, these mutant cells maintained high viability throughout the experiment (Figure 7B). For yKH063, there appeared to be an early VR starting at day 2, followed by another one at day 9 that lasted to day 20. The yXL171 parental strain committed a single VR starting at day 5. In the *aro8*Δ *aro9*Δ mutant culture, VR appeared to occur at a higher viability score than the parental control (compare the colony sizes of day 5 yXL171 and day 9 yKH063, Figure 7B), suggesting that increasing the endogenous supplies of aromatic alcohols caused cells to relax the threshold for the time to execute VR earlier, similar to the case of exogenous provision of such compounds (Figure 6). To test whether the elevated survivorship of these two strains was correlated with synchronized cell division, SMAS assays were conducted in parallel with the YPD spot assays. Coincidental to the ascending population viability was a boom in clonogenesis on the spent medium agarose slabs (arrows in Figure 7B; see Supplemental Figure 14 for the SMAS photos). Intriguingly, the emergence of these SMAS colonies corresponded to the “foothills” of the rising peaks of viability in the YPD spot assay (e.g., day 7 of yXL171, and days 9 and 11 of yKH063), suggesting that cell division was indeed an underlying mechanism for the increased population viability as revealed by the spot assay done on YPD plates. These genetic data therefore corroborated the notion that aromatic alcohols stimulate VRQT.

## Discussion

From the data presented above, we conclude that VRQT is an innate developmental program that extends population viability during starvation. This phenomenon is seen in haploid and diploid laboratory strains, in a commercial wine yeast, and in strains isolated from clinical specimens, woods, or vineyards. That post-VR clones repeat the same fluctuation of viability through the stationary phase without a clear extension of chronological lifespan argues against the acquisition of a spontaneous immortal or longevity mutation. The observations that diploid cells exhibit VRQT within a couple weeks are quite intriguing in that diploid cells have the options of sporulation to stay alive for a long period of time(Huang and Hull, 2017), and pseudohyphal growth to forage nutrients(Kumar, 2021). However, common laboratory protocols for sporulation call for the use of potassium acetate and sometimes a trace amount of raffinose(Elrod et al., 2009) when all other nutrients are kept to a minimum. Similarly, pseudohyphal growth is typically triggered by growing the amenable diploid strains on solid agar with low ammonium sulfate(Gimeno et al., 1992). In contrast to these two more restrictive growth conditions, VRQT happens in cells grown naturally old in a complete medium; there are no appreciable asci or pseudohyphae (Supplemental Figure 8). Thus, yeast cells have the capacity to engage different developmental pathways to deal with crises. Because VR is an age-dependent phenomenon, it remains to be delineated whether VRQT is driven by aging, starvation, or both.

Viability fluctuation of stationary-phase cultures has been noted previously(Fabrizio et al., 2004, Herker et al., 2004, Maqani et al., 2018), which was thought to result from adaptive regrowth mutations that allowed cells to grow on nutrients released from apoptotic cells. However, no evidence for cell division was shown in these prior studies. The presumptive adaptive regrowth “GASPing” mutations remain elusive. It is interesting to note that several mutations of *CYC8*, a multifunctional transcriptional co-repressor, rescued the shortened chronological lifespan of *snf1*Δ cells(Maqani et al., 2018). Though deemed as GASPing mutations, these *cyc8* mutants seemed to act as allele-specific suppressors for the *snf1*Δ mutant. Whether they are bona fide mutations that drive adaptive regrowth is unclear. As to whether apoptosis contributes to VRQT, three lines of evidence argue against this model. Firstly, deleting the metacaspase gene *YCA1*(Madeo et al., 2002) has no effect on the prevalence of VR (Table 1). Secondly, while the spent medium collected from a 7-day-old lean cell culture triggered VR of fat cells, the wildtype strain yMK839 at the same age did not respond to this conditioned medium (Figure 3). Nutrients, therefore, are not likely the main trigger for VRQT. Lastly, VR can be stimulated by supplementations of aromatic alcohols but not glucose or ammonium sulfate (Figures 5B and 6). We favor the hypothesis that VRQT, like other cellular growth and development programs, is built on a signaling network that includes inducers and receptors, and demands certain metabolic readiness. Metabolites such as indole acetic acid (Figure 6B and Supplemental Figure 13A) may act as an inhibitor of VRQT. The identification of the receptor and other components of this signaling pathways awaits further research.

While the separation of Q and NQ cells in stationary-phase cultures is well-documented(Allen et al., 2006), the heterogeneity among quiescent cells, such as the cell size and the replicative age, is also noted(Sagot and Laporte, 2019). Differences in the experimental setup, the age of the culture, and the means of inducing starvation further contribute to the challenge in isolating quiescent cells for the delineation of subcellular features key to the preservation of survival in starvation(Opalek et al., 2023). The SMAS assay developed in this work not only provides the definitive proof of robust cell division with scarce food and ample metabolic wastes, but also reveals a fascinating aspect of yeast cell population dynamics in aged cultures, that is, the viable cells in an aged culture can be divided into two categories. The first subpopulation is comprised of cells that adapt to the deprived and harsh conditions by eliciting the VRQT survival program to proactively form microcolonies in the same hostile environment. The second subpopulation conforms to the canonical quiescent cells that stay in the G^0^ phase until the environment improves. The coexistence of these two subpopulations highlights the resilience and adeptness of yeast. Furthermore, the behavior of the VR founding cells resembles that of adult stem cells in humans(Cheung and Rando, 2013) in that these yeast cells stay dormant but spring to action when perceiving the signals (e.g., aromatic alcohols), thereby exponentially supplying viable cells. If conditions are not improved, or the supply of the presumptive receptor subsides, cell division stops and the newly produced progeny loses the mitotic potential, as if entering senescence or becoming terminally differentiated. This concerted loss of mitotic potential contributes to the decline of the population viability following the VR-driven division, hence a wave of VR but not sustained viability. If the overall growth condition improves in time, these cells can quickly take over the niche and become a dominant clone among potential competing parties. Conceivably, technologies that can differentially isolate these two subpopulations for omics studies would shed light on the key elements supporting long-term survival.

The population density-dependent clonogenesis on spent medium led to the findings that aromatic alcohols stimulate VRQT (Figures 5 and 6). Our attempt to diminish these metabolites by deleting the *ARO8* and *ARO9* genes according to the Ehrlich pathway resulted in unexpected outcomes – instead of reducing the production, *aro8*Δ *aro9*Δ mutant cells accumulated higher levels of all three aromatic alcohols in the stationary phase (Figure 7A). This observation is in agreement with the findings by Chen and coworkers(Zhang et al., 2021), and suggests the existence of different pathways for aromatic alcohol biosynthesis. In the process of phenylalanine and tyrosine biosynthesis through the shikimate pathway(Braus, 1991), the intermediate prephenate can bypass the reaction catalyzed by the aromatic aminotransferases Aro8p and Aro9p in the Ehrlich pathway (see Supplemental Figure 15). In this case, prephenate is converted to phenylpyruvate and 4-hydroxyphenylpyruvate by Pha2p and Tyr1p, respectively(Braus, 1991). Phenylpyruvate and 4-hydroxyphenylpyruvate are then converted by Aro10p or an equivalent enzyme to the corresponding acetaldehyde for further reduction into alcohols. Because Aro8p and Aro9p are reversible aminotransferases, deleting these two enzymes can in theory reduce the loss of α-keto acid to amino acids. Coupled with the feeding of prephenate from the shikimate pathway, the production of PheOH and TyrOH can be increased in *aro8*Δ *aro9*Δ background. On the other hand, Aro8p and Aro9p are the only enzymes known to catalyze the formation of indole-3-pyruvate, the “Trp equivalent” of phenylpyruvate. It is unclear as to why the TrpOH level was also increased in *aro8*Δ *aro9*Δ cells (Figure 7A). We suspect that a new pathway specific to the non-fermentative growth condition for VRQT is responsible for tryptophol synthesis. Our preliminary genetic mutant screens have not found a gene related to aromatic amino acid metabolism such as *ARO10*, *BAT2*, *BNA3*, or *YER152C* (a homologue of *ARO8* and *ARO9*) to be important for VRQT (Table 1). More comprehensive tests are needed to construct the pathway and the regulation of this survival program.

Quiescence is the predominant state of most cells in the wild and in multicellular organisms(Sun and Gresham, 2021, Cheung and Rando, 2013, O’Farrell, 2011, Coller, 2011). In yeast, there is a colossal set of changes associated with quiescence for long-term survival. Significant alterations in the cell wall structure, the lipidome, transcriptome, metabolome, and proteome indicate sophisticated coordination in functions of different organelles and chromatin structures(Laporte et al., 2016, Miles et al., 2013, Brauer et al., 2008, Bradley et al., 2009, Schroeder and Shadel, 2014, Klose et al., 2012). In the stationary phase, reactive oxygen species from normal cellular metabolism and environmental irradiation may damage biomolecules, including lipids, proteins, RNAs, and DNA(Gangloff and Arcangioli, 2017, Li et al., 2017, Jones, 2015, Lu and Finkel, 2008, Chen et al., 2007). Over time, damage that escapes the plethora of cellular repair and scavenging mechanisms may be passed on to the progeny cells through VRQT. However, these damages, except DNA mutations, are not inheritable. It is formally possible that a fraction of such DNA mutations may give the offspring cells better competitiveness and drive evolution. In our opinion, a GASP-like mutation must meet two criteria. Firstly, it triggers efficient cell division. Secondly, it gives the progeny a sustainable growth advantage. Such mutations therefore should be rare. We have not detected growth advantage of any of the post-VR clones in competition experiments (e.g., Figure 2). We suspect that VRQT can potentially drive evolution by passing existing mutations to progeny, but VRQT per se, though formally possible, is unlikely to be driven by a mutation.

Mechanistically, the chronological lifespan is viewed as a tug-of-war between quiescent (Q) and non-quiescent (NQ) cells during starvation(Werner-Washburne et al., 2012). However, the pervasiveness of VRQT among strains of *S. cerevisiae* suggests that cellular division in a famine may be an important contributor to population longevity. We posit that in an aging, stationary-phase culture, the combination of the loss of clonogenesis (i.e., senescence and death), the successful establishment and maintenance of quiescence, and the expansion of a mitotically capable population from VR together determine the overall survivorship at the time of sampling. Frequent “mini-VR” activities, albeit brief, may bestow vitality on the population, therefore leveling the trend of time-dependent loss of the ability to divide. For example, several wild strains, such as the wine yeast yKH070, a clinical isolate yKH076, and yKH079 from oak exudate, exhibited 2 or 3 stabilization periods that lasted four or more days (arrows, Supplemental Figure 3). It is possible that a moderate number of cell division resupplies viable cells to “cancel” out the number of dying cells, thus interrupting the downward trend of survivorship. In other words, population longevity can be achieved by either delaying the aging of individual community members, or by revving up the resupply of fertile descendants. In the sector of infectious disease, this behavior may fortify the development of biofilms and the production of virulence factors(Zara et al., 2020, Albuquerque and Casadevall, 2012, Wuyts et al., 2018), so it is worth deeper scrutiny of the molecular mechanisms and their control of VRQT.

## Acknowledgments

We thank Gerald Fink for the generous supply of Σ1278b strain, and Eric Hegg and Zhen Fang for the advice and instrumentation of the quantification of aromatic alcohols with HPLC. The authors are in debt to Lee Kroos, Erich Grotewold, and Tommy Vo for critiques and suggestions on the manuscript. The authors also thank Nakoa Po, Kuang-Wei Wang, and Hsiao-Tien Chien Hagor and members in the Kuo lab for frequent discussion and feedback along the development of this project. This work was supported by funding from NSF (CMB1817324) to MHK.

## Author contributions

KCH made the observation of VRQT and performed most of the experiments. HYL quantified the aromatic alcohols in medium and performed sporulation experiments, and helped conduct AUC calculation. MHK designed the original SMAS assay. TRH helped design the knockout experiments. MHK and KCH wrote the manuscript together.

## Materials and methods

### Strains and medium

Yeast strains used in this study are shown in Table 1. YPD medium contained 2% glucose (Sigma-Aldrich), 2% peptone (BD Difco), and 1% yeast extract (BD Difco). SC medium (synthetic complete) contained 2% glucose, 5 g/l ammonium sulfate (Sigma-Aldrich), 1.7 g/l yeast nitrogen base without amino acids or ammonium sulfate (BD Difco), and complete amino acids as descried in(Sherman, 1991). Auxotrophic nutrients were supplied at four-fold excess as recommended(Hughes et al., 2012). 2-phenylethanol, tyrosol, and tryptophol (all from Selleck Chemical; Houston, TX) were added from 100 mM stocks in ethanol to SC medium and agarose slabs to achieve final concentrations of 10 μM or 100 μM. PSP2 medium had the following composition: potassium phthalate 8.3 g/L, yeast extract 1 g/L, 1.7 g/L yeast nitrogen base without amino acids or ammonium sulfate, ammonium sulfate 5 g/L, potassium acetate 10 g/L, pH 5. Sporulation medium contained potassium acetate (3 g/L) and raffinose (0.2 g/L).

### Yeast methods

Yeast cells were transformed using the lithium acetate method(Gietz et al., 1992). To create certain deletion strains, PCR reactions were performed using primers shown in Table 2. The *URA3* and *TRP1* genes of *K. lactis*, amplified from the pBS1539 and pBS1479 plasmids respectively(Markgraf et al., 2014), were employed as selection markers. The PCR products were purified using gel extraction for yeast transformation. Genomic insertion was verified by genomic PCR.

**Table 2.**
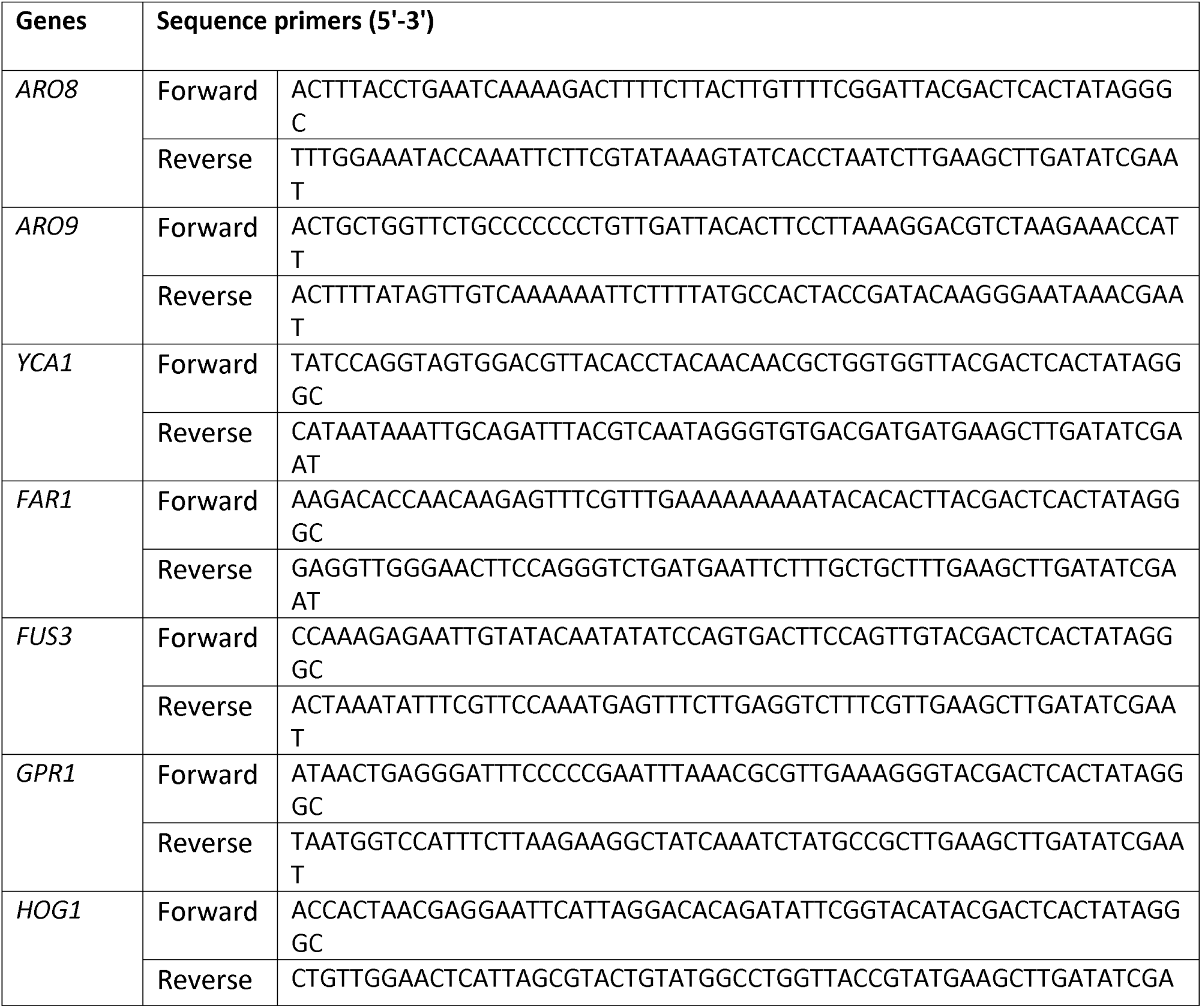
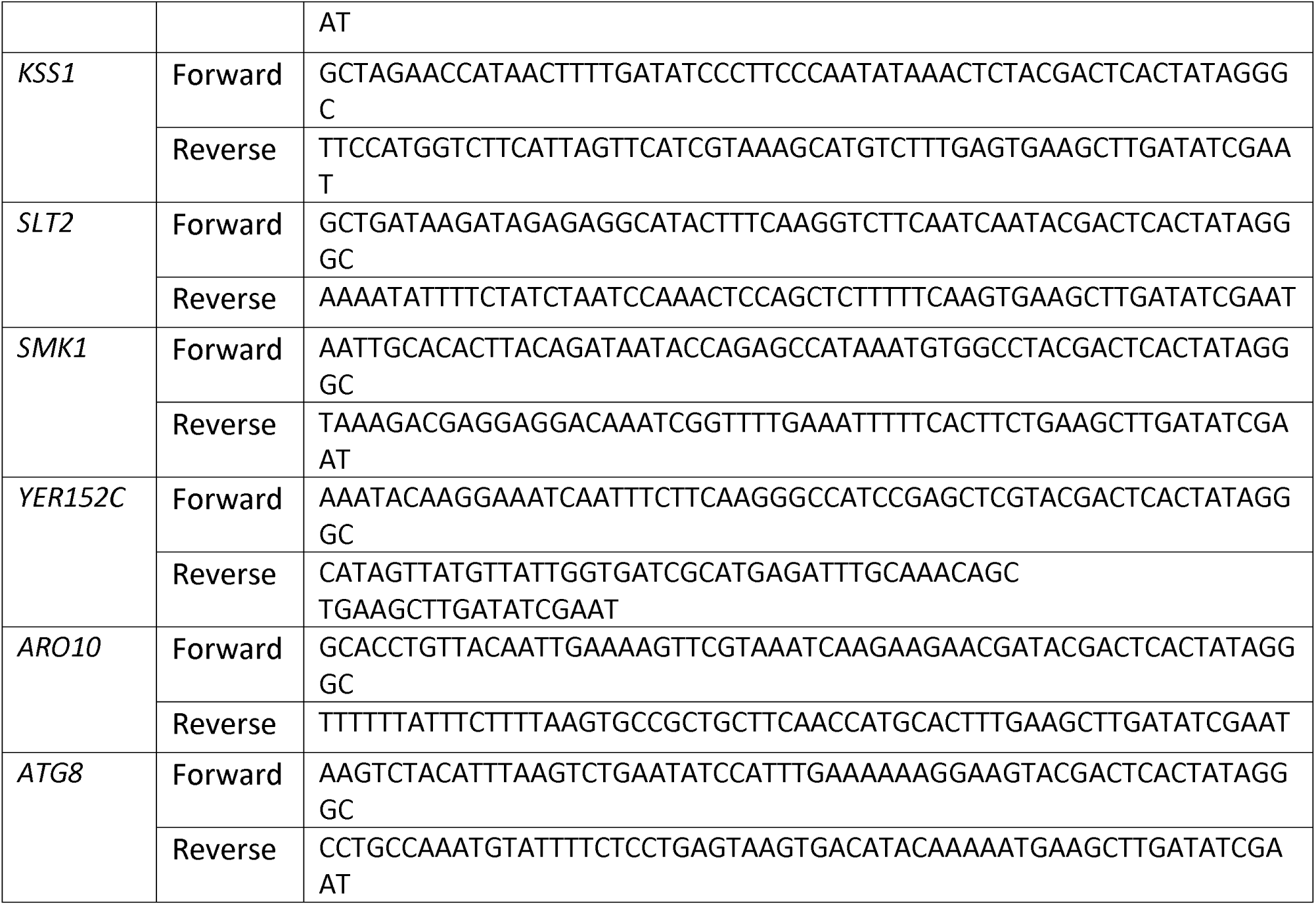
Primer sequences.

### Spot assay and viability scoring

Cells were cultured in synthetic complete (SC) medium at 37°C, using a rotating wheel (Thermal Scientific, Cel-Grow Tissue Culture Rotator) set to 60 rpm with a 13-degree tilt to ensure uniform growth. The cultures were grown in 50 mL tubes covered with aluminum foil for aeration, and each tube contained 25 mL of SC medium to ensure adequate aeration. At predetermined timepoints, the optical density at 600 nm (O.D._600_) was determined using a spectrophotometer. Following measurement, 1 O.D.^600^ cells were centrifuged at 14,000 rpm for one minute to pellet the cells at room temperature. The supernatant was decanted, and the cell pellets were resuspended in 100 μL of double-distilled water (ddH^2^O). A series of tenfold serial dilutions was prepared from this suspension. 5 μL from each dilution was spotted onto a YPD agar plate. These plates were incubated at 30°C for two days before colony growth was assessed. The growth score was quantified using a spot assay-based scoring system. Spots displaying 3-10 colonies were assigned 0.5 points, while those with more than 10 colonies received 1 point. The final growth score for each sample was calculated by summing the scores from all six spots of the dilution series. This resulted in a scale ranging from 0 to 6, where higher scores indicated higher viability. The Area Under Curve (AUC) was calculated using GraphPad Prism software (version 10, GraphPad Software, San Diego, CA), based on the growth scores of the experimental samples collected over a one-month period.

### Spent-medium agarose slab (SMAS) assay

1.5 mL of yeast cell culture was collected at the indicated time point and centrifuged in a microcentrifuge at 14,000 rpm for 1 minute at room temperature. The supernatant was transferred to a 2 mL Eppendorf tube with 0.03 g of agarose to make a final concentration of 2% (w/v). The mixture was heated in a boiling water bath for 30 minutes to melt the agarose, and poured into a 12-well plate, 1.5 mL per well. The agarose slabs were allowed to solidify at room temperature and the plates were stored at 4°C before use. For cell inoculation, each solidified agarose well was spotted with 3x dilution from 0.5 O.D._600_ cells (∼10,000,000 cells) in a volume of 5 μL of water suspension. The cells were incubated at 37°C for two or three days to promote initial colony formation. Subsequently, the agarose slabs with yeast inocula were excised and transferred onto YPD agar plates and incubated at 30°C for an additional two days to assess the overall viability.

To record the growth of each inoculum on SMAS or the subsequent YPD transplantation, pictures were taken one inoculum at a time under an Olympus SZ60 stereomicroscope (10 x eyepiece, 1 - 6.3 x objective). Typical images were taken at 25 – 30 x magnification with an iPhone 7. This magnification captures the entire inoculum (5 µL initial volume, approximately 5 mm in diameter) with individual cells clearly discernible.

### Medium treatment

2 mL of 5-day-old yXL171 media were separated from cells by centrifugation (14,000 rpm for 1 minute at room temperature). The cell-free media were then treated with either 6 units of DNase I (Invitrogen), 5 units of RNase A (Sigma), or heated in a boiling water bath for 30 minutes. These treated media were used for medium swap experiments at indicated time points.

### Aromatic alcohol quantification by HPLC

The quantification of aromatic alcohols in yeast medium was described previously(Zupan et al., 2013). Briefly, yeast medium was collected at predetermined time by centrifuging at 10,000 rpm for 5 minutes at 4°C and the supernatant was further filtered through a 0.2 µm filter. The cleared medium was stored at 4°C until analysis. High-performance liquid chromatography (HPLC) analysis was performed on an Agilent 1260 Infinity system equipped with a diode array detector. The reverse phase column (Ascentis Express C18, 2.7 μm, 4.6 mm × 150 mm) was used with isocratic mobile phase 80:20 (v/v) H^2^O/acetonitrile. The column temperature was set at 30°C. Chromatographic separation was achieved by isocratic elution at a flow rate of 0.5 mL/min. The total run time was 18 min, and the injection volume was 50 µL. The three aromatic alcohols, 2-phenylethanol, tryptophol, and tyrosol were detected at their optimal wavelengths of 255, 225, and 225 nm, respectively, using the purchased compounds as the standard.

### Sporulation

Sporulation was conducted according to (Kassir and Simchen, 1991). 2 O.D.^600^ of overnight diploid strain culture was inoculated to 2 mL of PSP2 medium and shaken at 250 rpm at 30°C to achieve good aeration. When the cell concentration reached 1 O.D.^600^/mL, 1.5 mL cells were harvested by centrifugation at 5,000 rpm for 5 minutes, washed once with sterile water, and re-suspended in 1 mL SPM medium. Asci can be seen after two days of incubation at 30°C with 250-rpm shaking. Cells were strained with DAPI (4,6-diamidino-2-phenylindole) in the mounting medium (1 mg/mL p-phenylenediamine, 90% glycerol, 0.1 μg/mL DAPI) to visualize the the nuclei of spores or non-sporulating cells.

## Supplemental figure titles and legends

**Supplemental Figure 1. VRQT also occurs at 30°C.**

yMK839 (wt), yXL171 (lean) and yXL006 (fat) strains were grown to the stationary phase in SC medium at 30°C, and sampled for YPD spot assays to compare the viability. Shown are representative results from three biological repeats.

**Supplemental Figure 2. Among laboratory yeast strains tested, only W303 cells lack VRQT at 37°C.**

Five laboratory strains were tested their ability to perform VRQT at 37°C. The survival scoring chart is shown on the right. Only W303 failed to show VRQT.

**Supplemental Figure 3. Many yeast wild strains perform VRQT.**

Prototrophic wild strains isolated from different sources (see Main Text for details) were subjected to VRQT tests at 37°C. In addition to VR, many wild strains showed significant stabilization of the viability (arrows). These are likely due to “mini-VR” that supplied sufficient viable cells to even out the net loss of viability.

**Supplemental Figure 4. Aged W303 cells showed VRQT when cultured at 30°.**

The ability to perform VRQT by W303 cells was tested at 30°C. A VR was recorded between days 42 to 48. When day 42 culture was aliquoted and transferred to the day 5 spent medium from the lean cells (yXL171), W303 cells grew better. However, when an identical medium swap aliquot was grown at 37°C, the growth was completely inhibited, suggesting that W303 cells were defective in coping with sustained growth at 37°C.

**Supplemental Figure 5. Spot assay images of the three pre-VR yXL171 tests used for AUC comparison with post-VR clones.**

**Supplemental Figure 6. Medium swap can change the course of viability resurgence.**

Shown are the complete set of experimental results for Figure 3.

**Supplemental Figure 7. VRQT inducers resist (A) RNAse, DNAse, and (B) boiling.**

**Supplemental Figure 8. Diploid cells go through VRQT without eliciting sporulation.**

A. Viability changes of three diploid strains. B. DAPI staining to show nucleus of each cell sampled at day 23. Asci are manifested by the presence of 4 nuclei (red arrows), which were only seen when cells were cultivated in sporulation medium (SPM).

**Supplemental Figure 9. Three additional examples of population density-dependent microcolony formation on SMAS.**

**Supplemental Figure 10. After 2 days of incubation on an agarose slab, the colony achieves 15 generations with a size of around 500 μm in diameter.**

Colonies of ∼0.5 mm in diameter were plucked and suspended in water for cell counting in a hemocytometer. The size and number of cells from each colony are shown in the table.

**Supplemental Figure 11. Formation of colonies on spent medium coincided with the viability resurgence on YPD.**

The scores of the spot assay and the SMAS assay were presented as 2-D area plots. Blue and orange areas represent the YPD spot assay and the SMAS assay, respectively. Note: SMAS assays were done on day 5 – day 13 cultures only.

**Supplemental Figure 12. Tryptophol is the most potent VRQT inducer among the three aromatic alcohols.**

The zoom-in images of the rightmost inoculum of the 10 µM tryptophol treatment slabs are shown on the right (* and **). Several microcolonies on SMAS are indicated by arrows. Note that the YPD growth was conducted for only 18 hours to allow the SMAS microcolonies to grow bigger but the secondary YPD-only colonies were still small.

**Supplemental Figure 13. Indole-6-acetoc acid (IAA) inhibits VRQT but not the log-phase cell growth.**

(A) Stationary-phase lean cells were 3-fold serially diluted and spotted to SC complete medium supplemented with 100 µM of indole-6-acetic-acid (IAA). The plates were then incubated for 24 hours at 37°C. IAA completely inhibited VR with zero colony observed (see Main text Figure 6). (B) 0.5 O.D._600_ log-phase yXL171, yMK839, and yXL006 cells were spotted to fresh SC medium supplemented with either 100 µM tryptophol (TrpOH) or 100 µM indole-6-acetic-acid (IAA). The plates were then incubated for 24 hours at 37°C. All inocula were scored as “lawn”. Neither compounds caused differences in cell growth.

**Supplemental Figure 14. Cells lacking *ARO8* and *ARO9* genes exhibit efficient VR and a growth advantage. Only 107 cells were spotted for assessment.**

Shown are SMAS assay results. The number of colonies are shown in the table.

**Supplemental Figure 15. Abbreviated schematics of the Ehrlich and shikimate pathways for aromatic alcohol metabolism.**

This figure is adapted from Hazelwood et al. (2008), Cordente et al. (2019), and Gonzalez-Ramirez et al. (2024)

