## supplemental figures for "Coordinated Division of Selective Stationary-Phase Yeast Cells Expands the Population Survivorship in Quiescence"

Hsiao et al.,

**Summary**

Budding yeast employs a variety of survival strategies in response to starvation, including pseudohyphal development, invasive growth, and sporulation. Here we report an innate pathway of "viability resurgence in quiescent time" (VRQT) by aged cultures to preserve and expand the population survivorship. Without additional nutrients, a few stationary-phase cells synchronously enter mitosis, increasing the population viability that does not likely result from mutations. VRQT is a population density-dependent collective behavior that can be triggered by quorum sensing aromatic alcohols. Genetic analyses demonstrate that VRQT is independent of many canonical pathways for cell growth, development, or stress responses. This community survival program allows yeast to proactively extend vitality amidst a common nutritional crisis. Wild strains isolated from clinical samples exhibit VRQT, suggesting that this survival mechanism must be considered when treating human infections.

#### Supplemental Figure 1

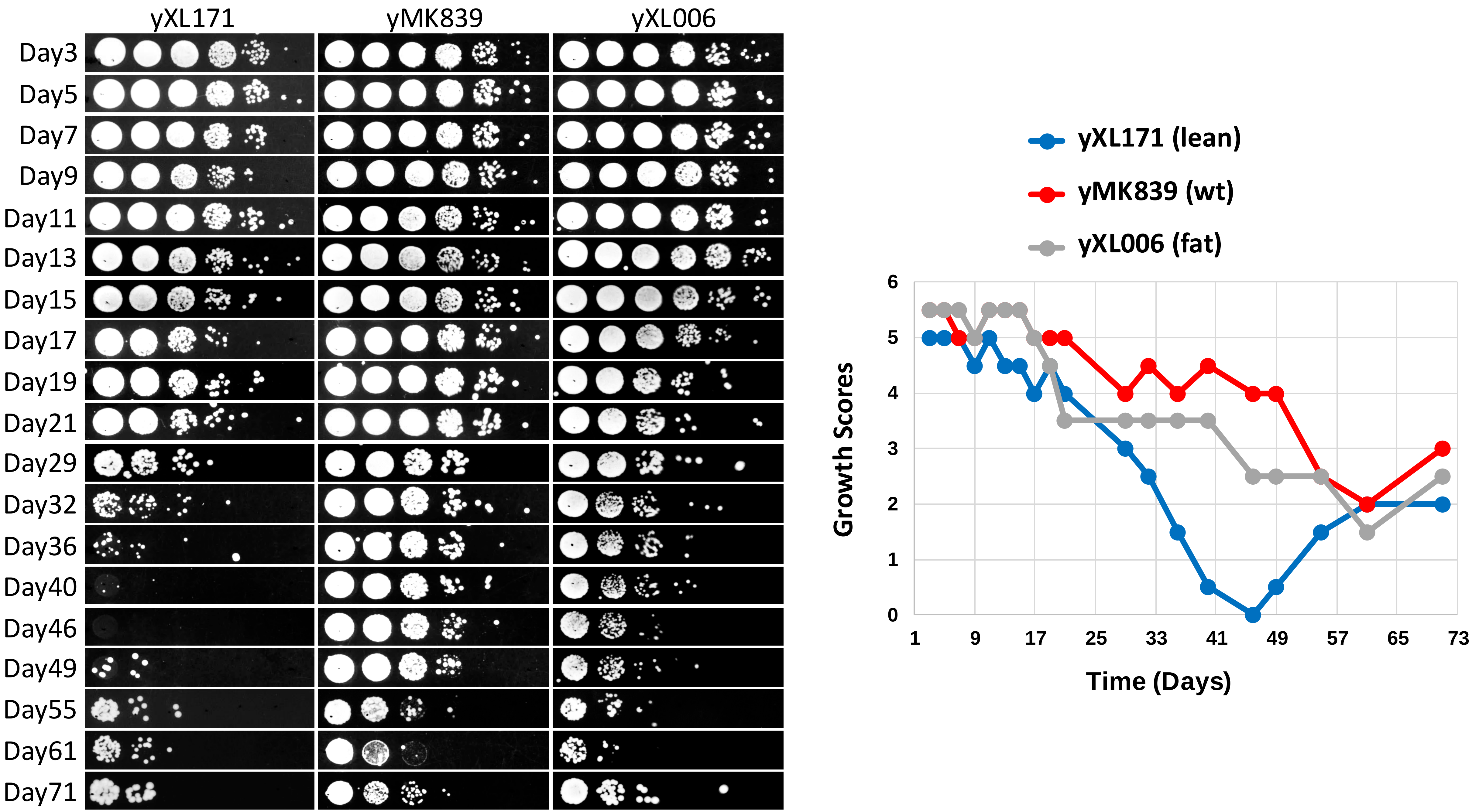

#### Supplemental Figure 2

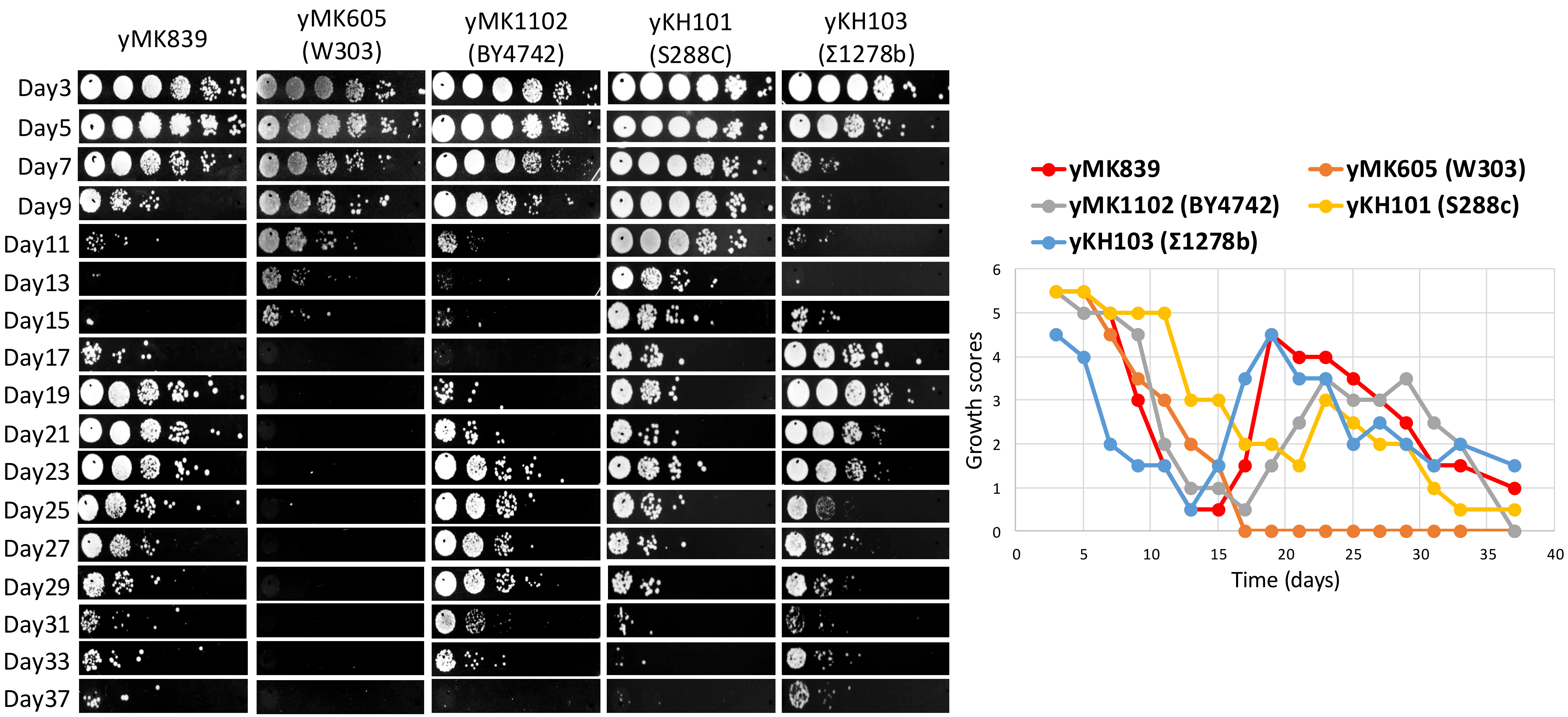

### Supplemental Figure 3

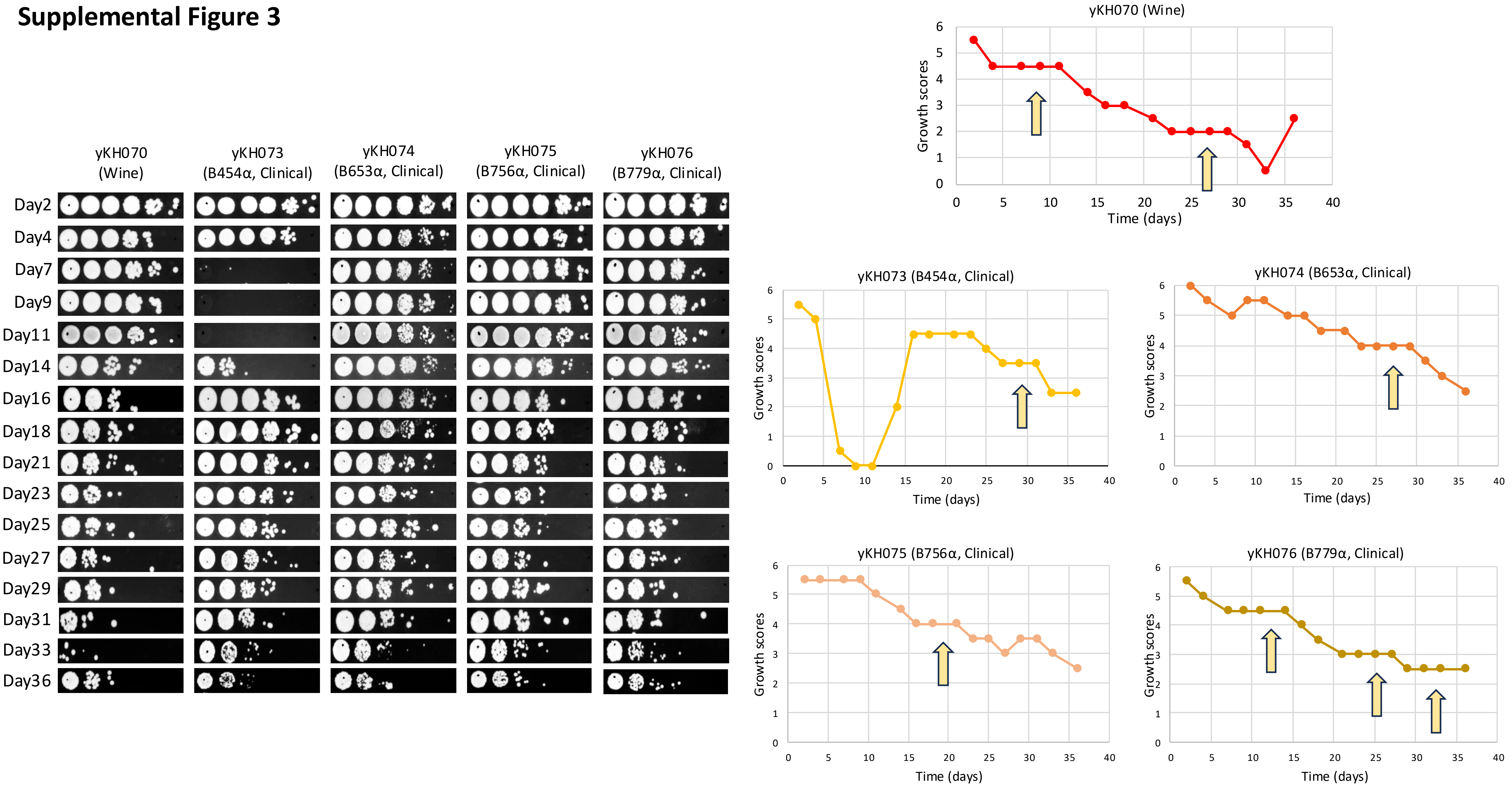

### Supplemental Figure 3 (cont'd)

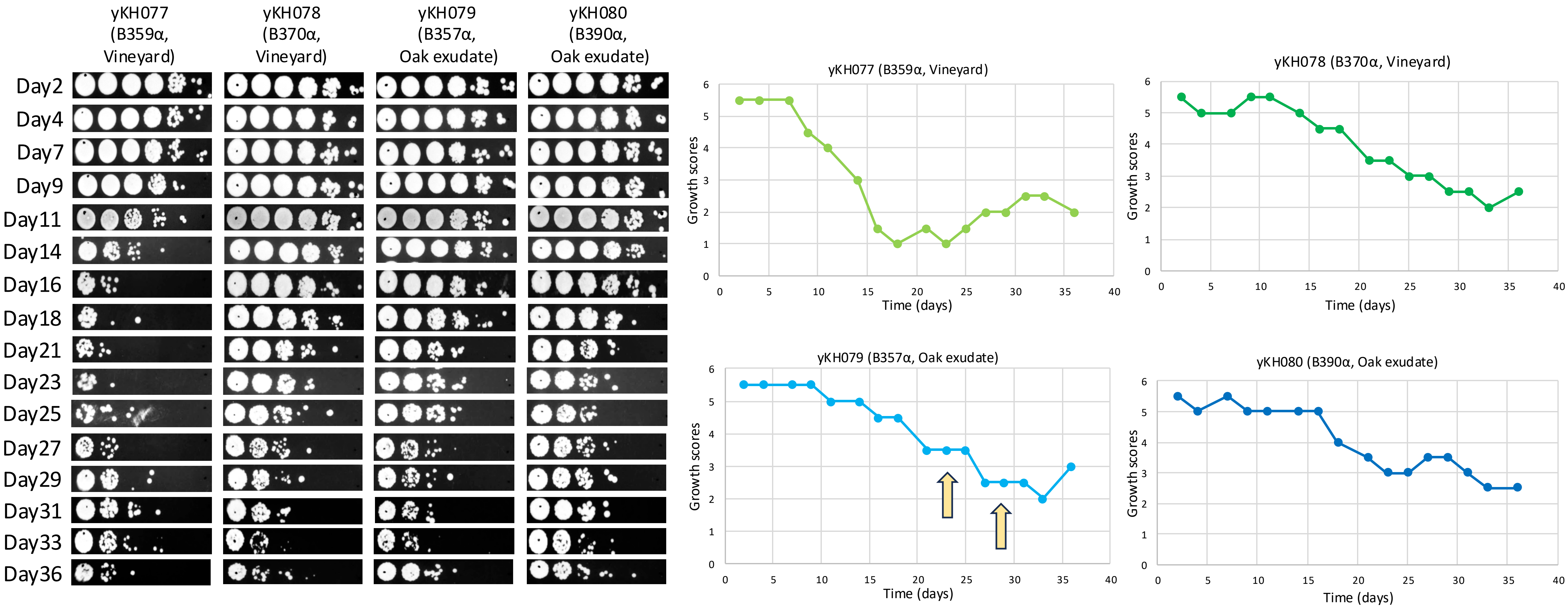

**Supplemental Figure 4**

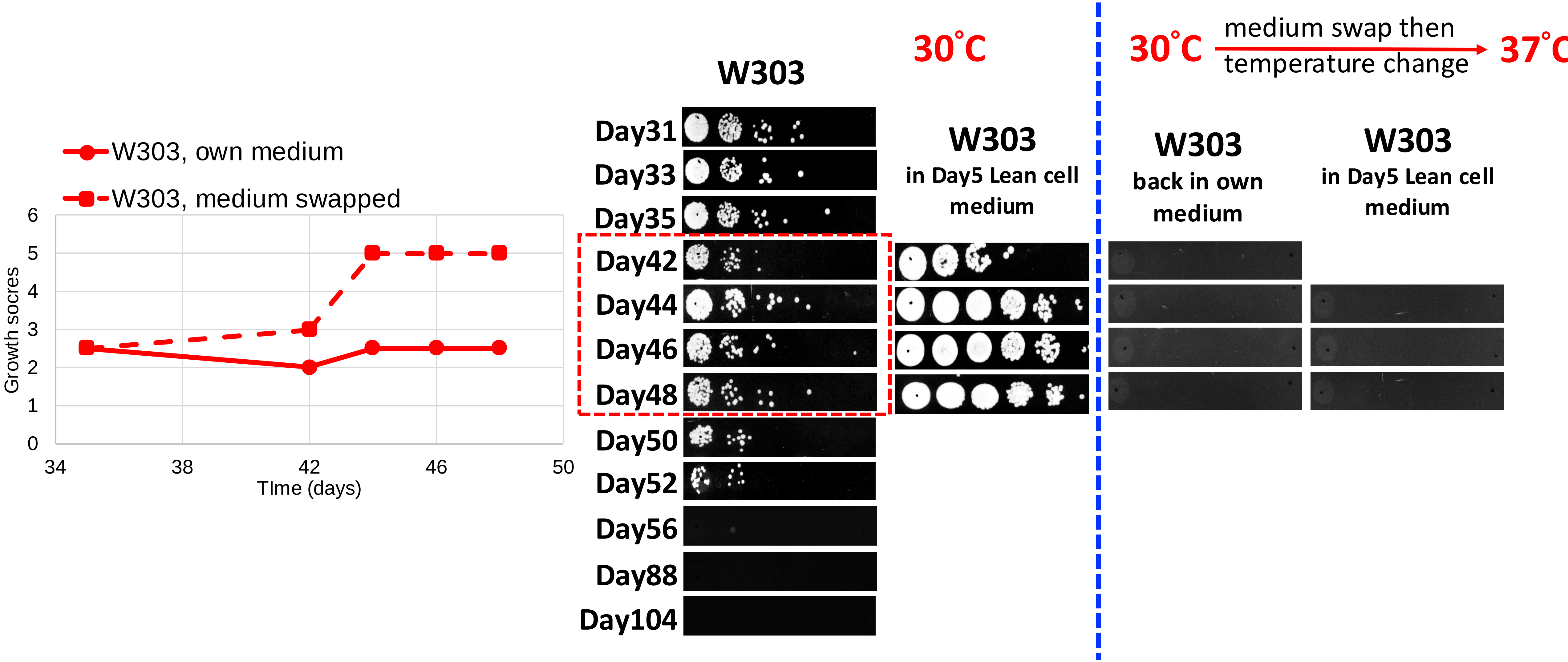

#### Supplemental Figure 5

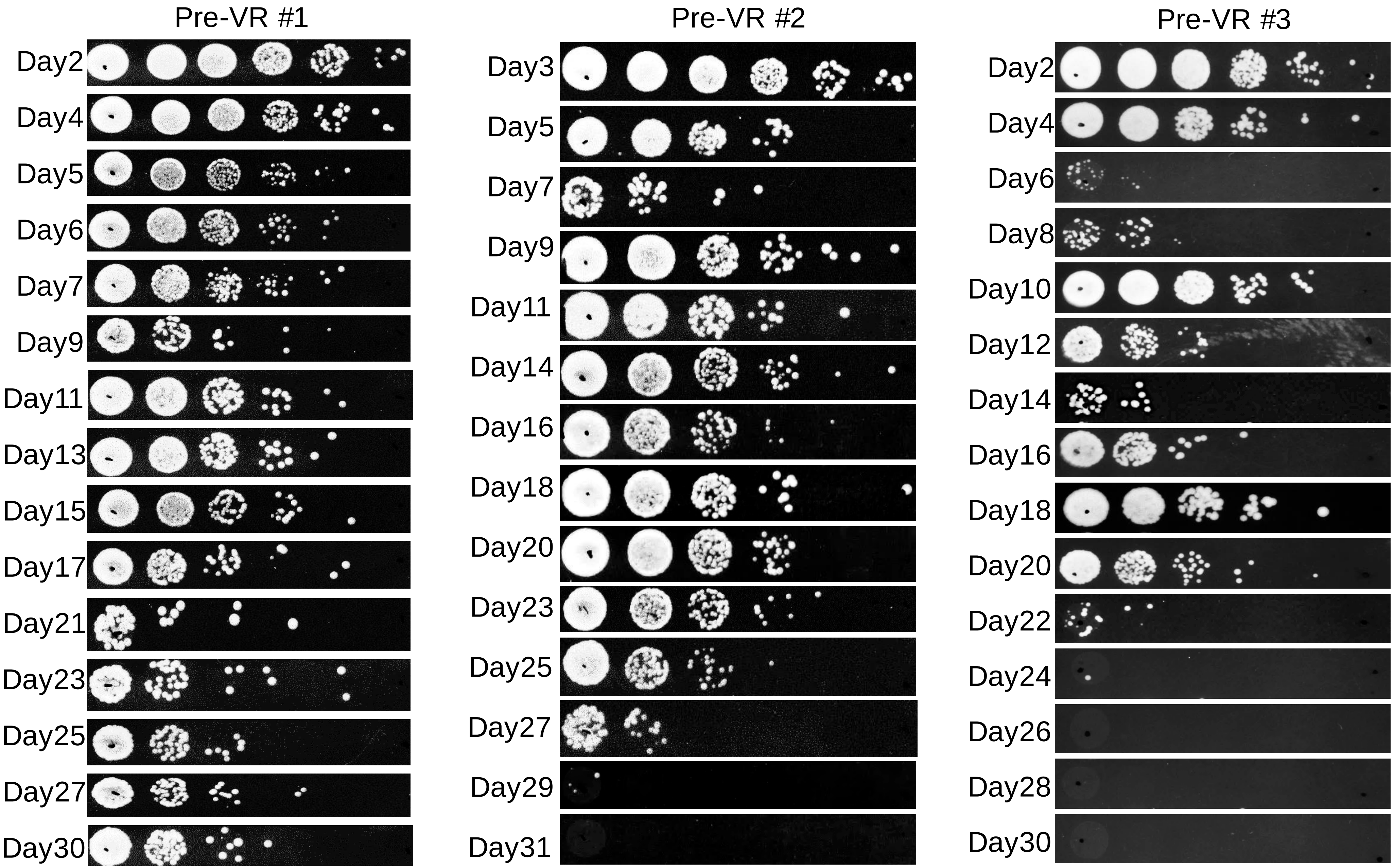

**Supplemental Figure 6**

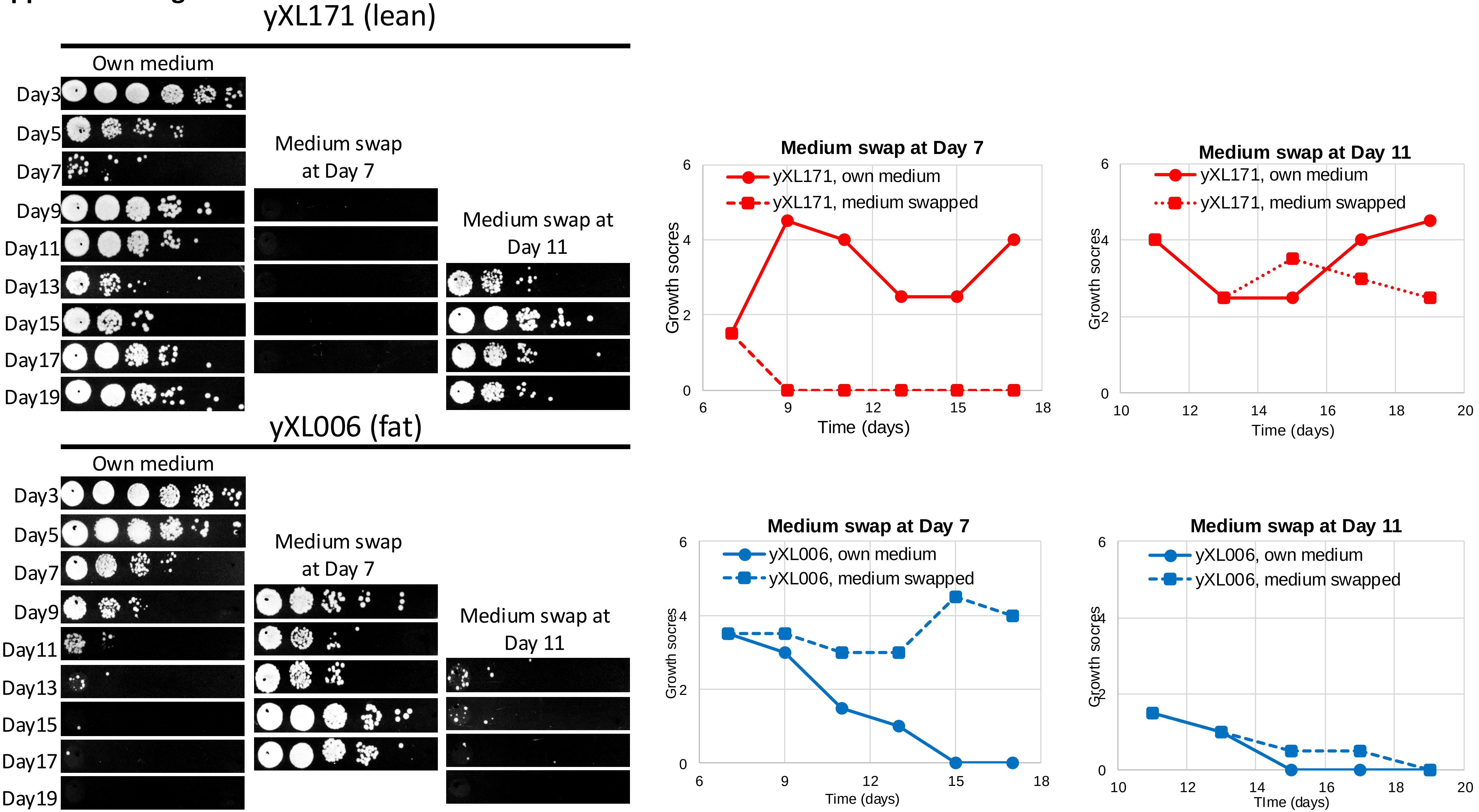

**Supplemental Figure 7**

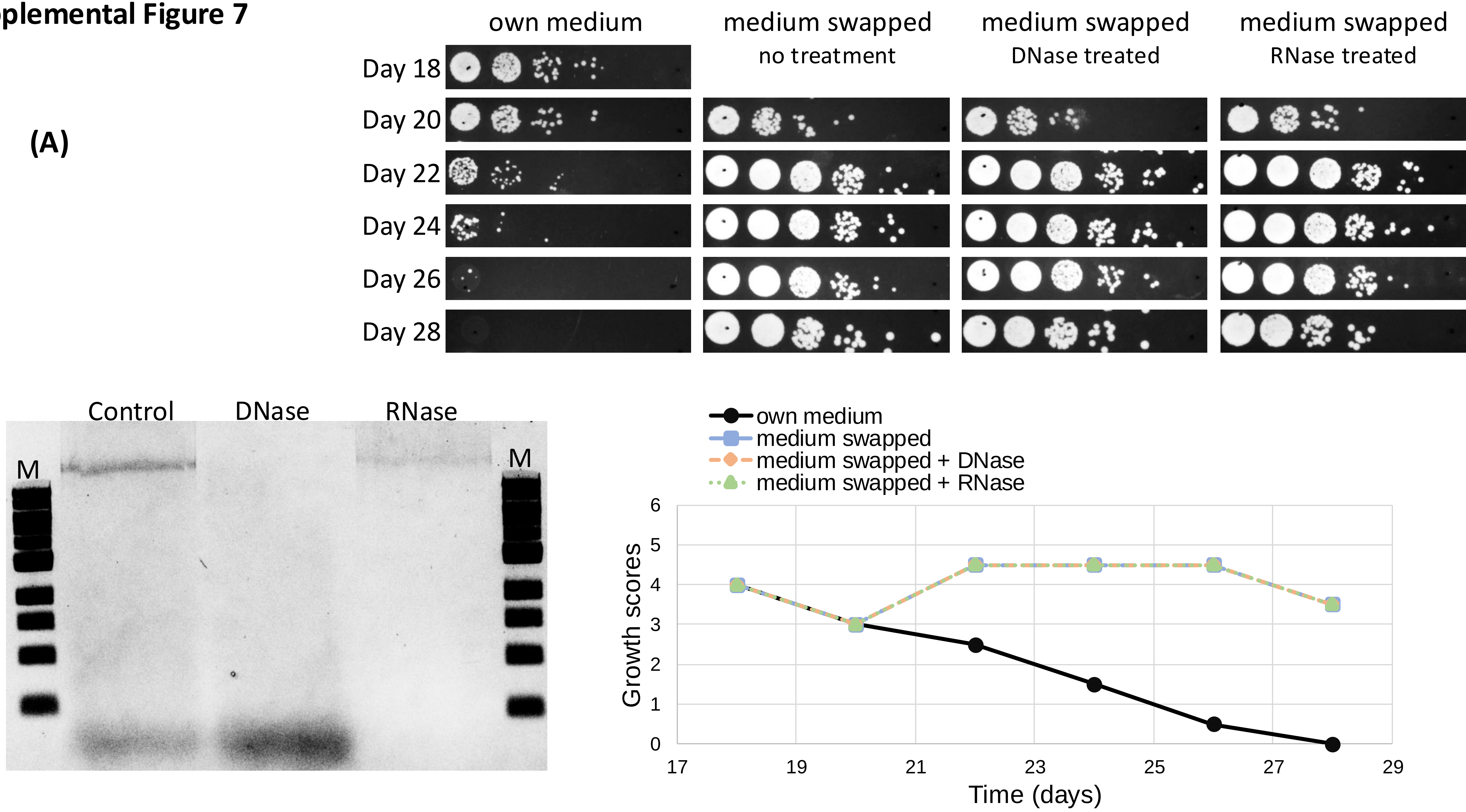

#### Supplemental Figure 7

### (B)

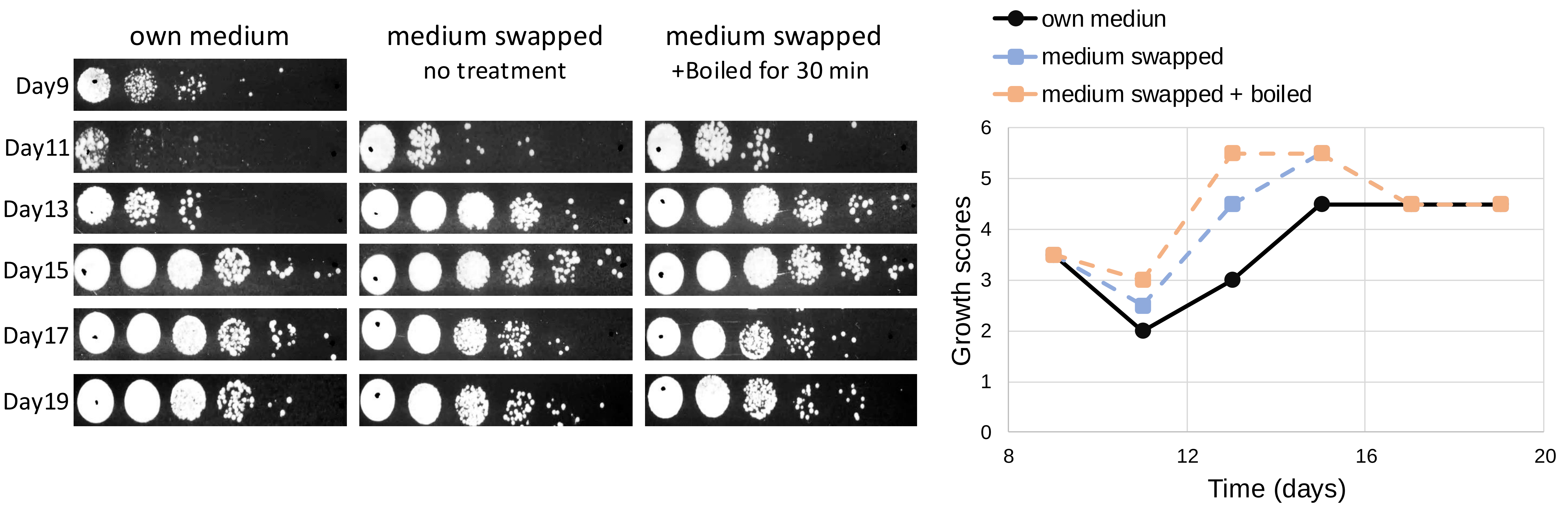

**Supplemental Figure 8**

Lean: yHL013 (*dga1Δ/Δ lro1Δ/Δ*)
WT: EJ72 (*DGA1+/+ LRO1+/+ TGL3 +/+)*
Fat: yWH074 (*tgl3Δ/Δ*)

**(A)**

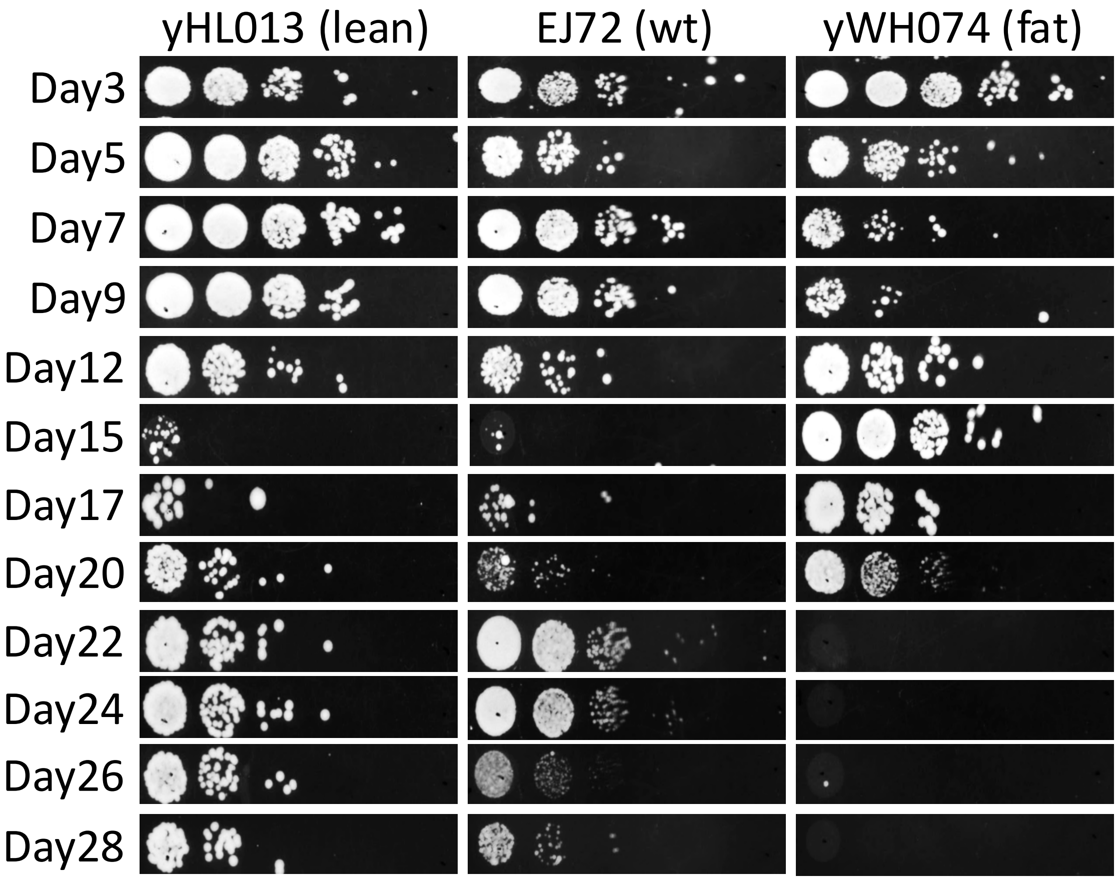


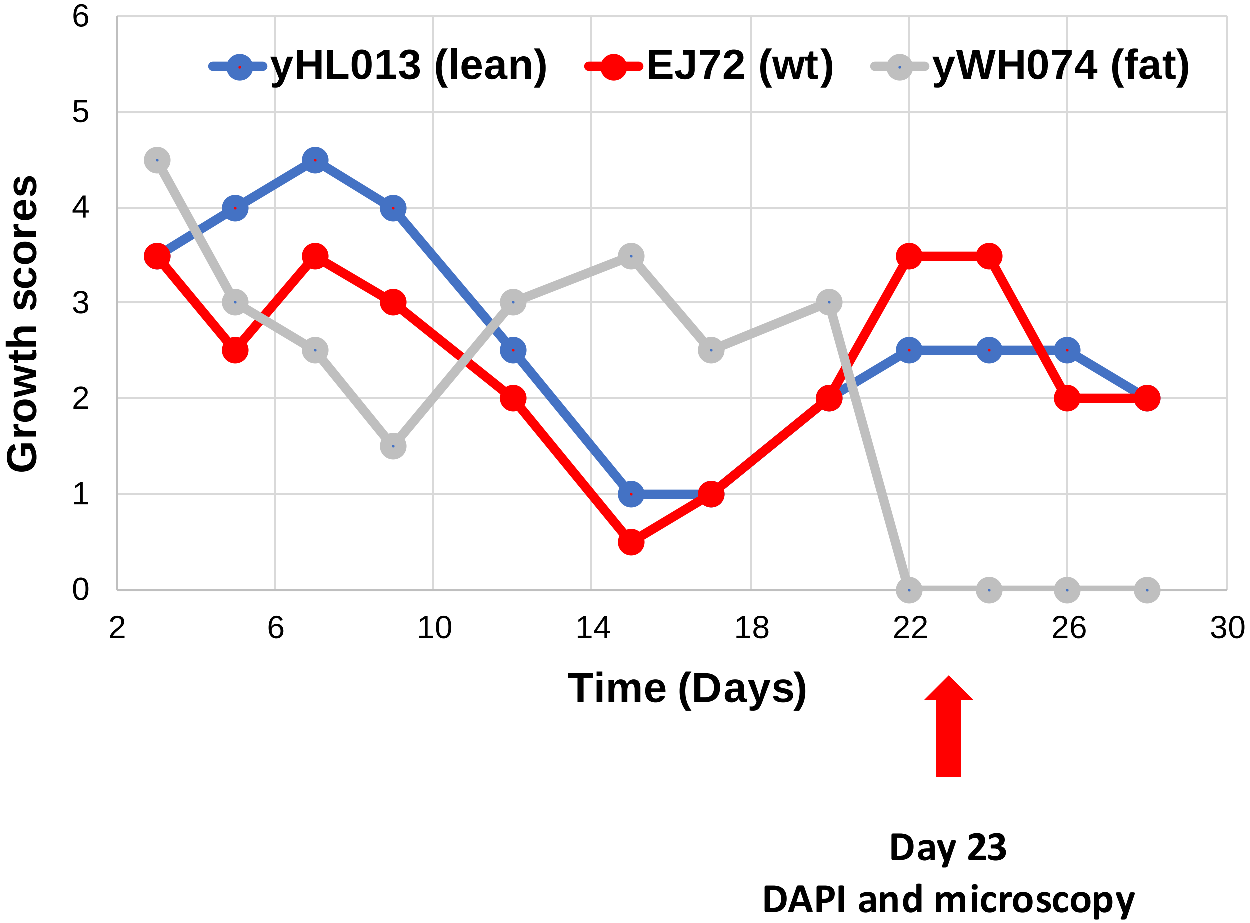

#### Supplemental Figure 8

**(B)**

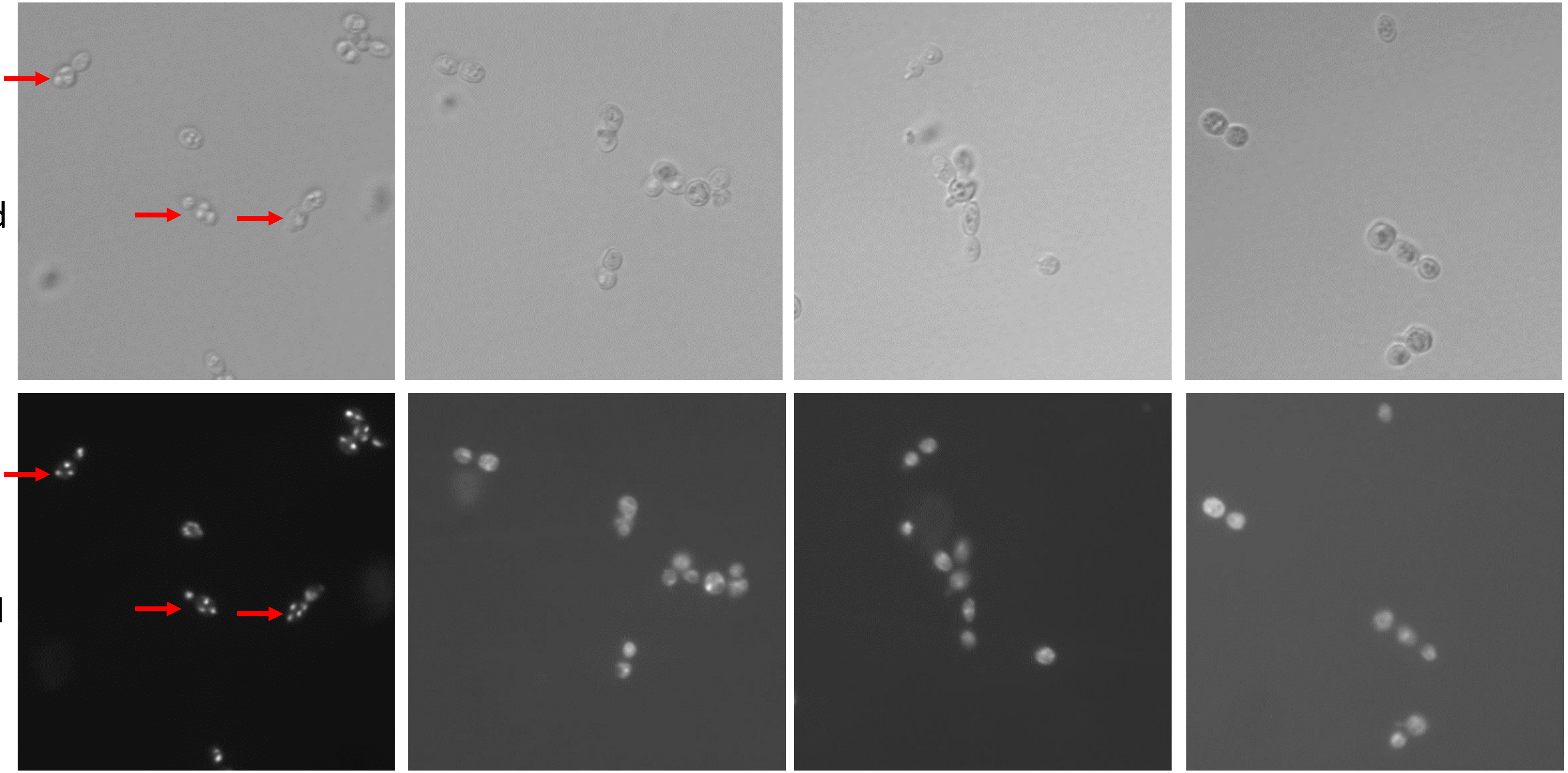

**Supplemental Figure 9**

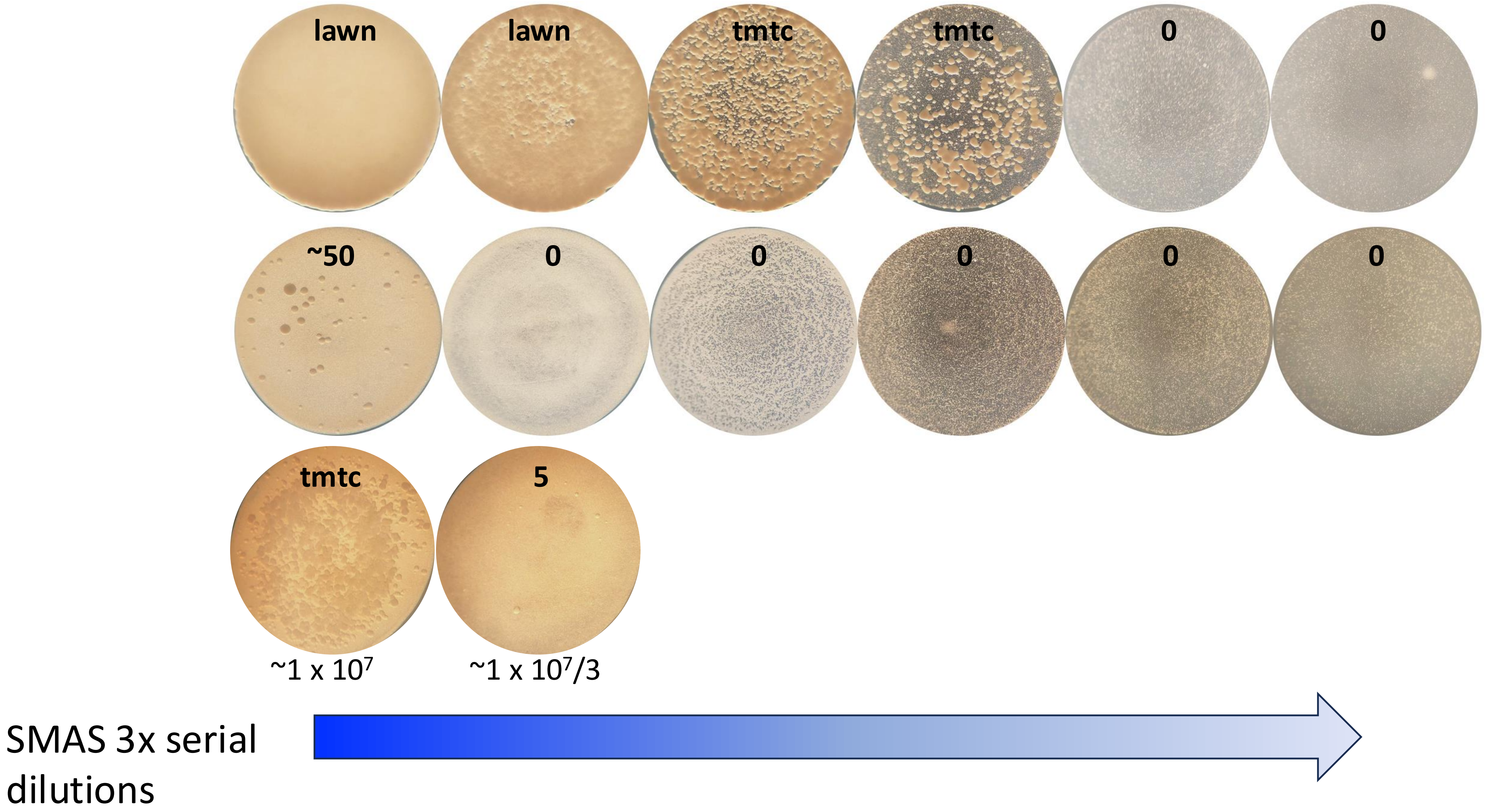

**Supplemental Figure 10.**

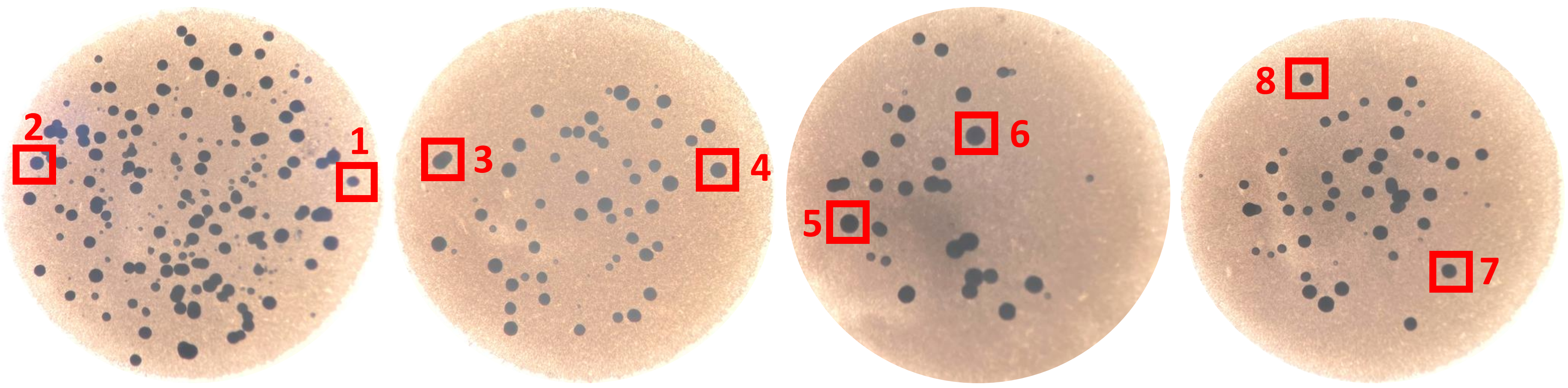


| | Colony_1 | Colony_2 | Colony_3 | Colony_4 | Colony_5 | Colony_6 | Colony_7 | Colony_8 | Average |
|---|---|---|---|---|---|---|---|---|---|
| Colony diameter (μm) | 461 | 489 | 711 | 524 | 560 | 514 | 512 | 451 | 527.75 |
| Cell number | 40250 | 19500 | 53750 | 45875 | 60625 | 44375 | 38125 | 45625 | 43515.62 |
| Generations | 15.29 | 14.25 | 15.71 | 15.48 | 15.88 | 15.43 | 15.21 | 15.47 | 15.34 |

**Supplemental Figure 11:**

(A)

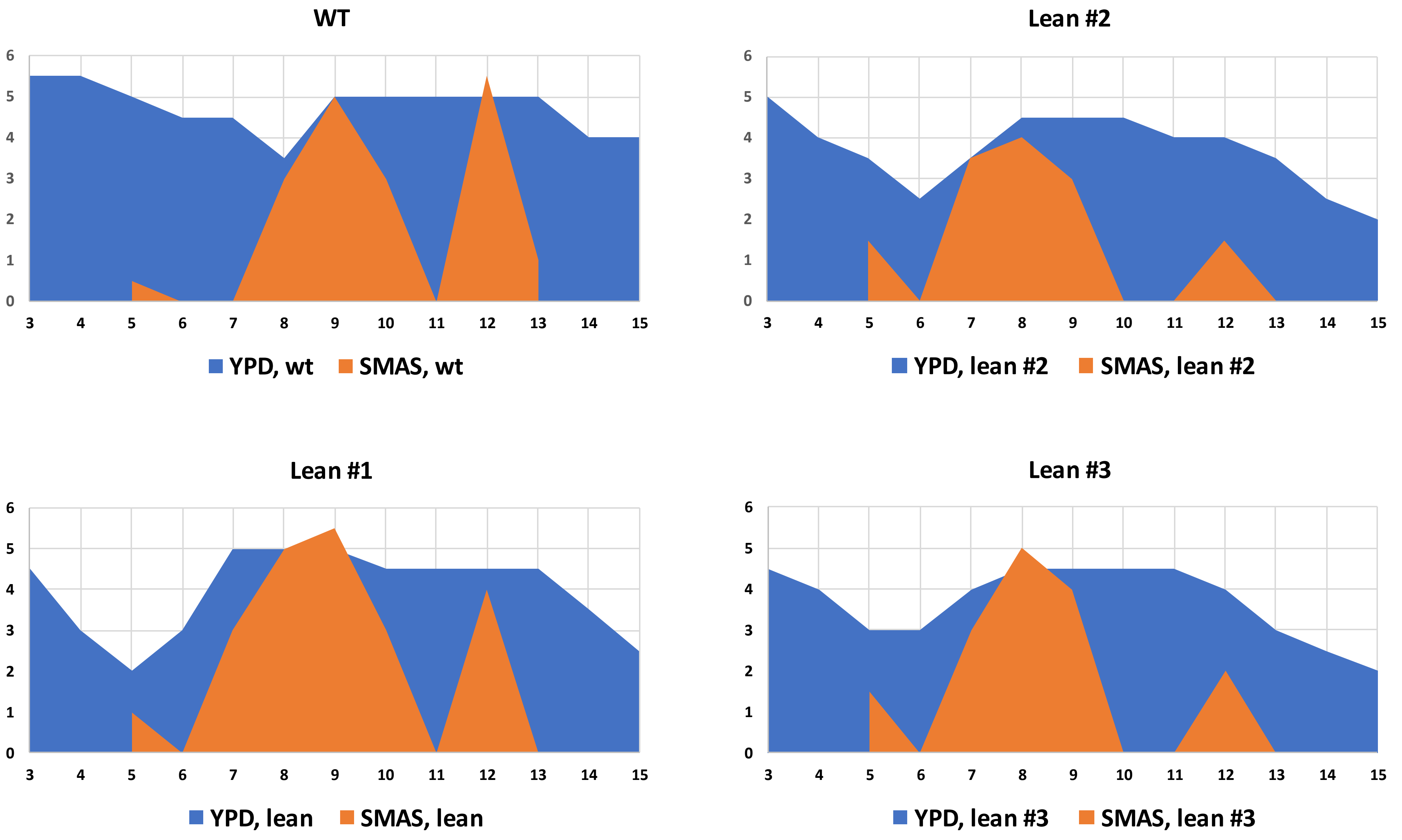

**Supplemental Figure 11**

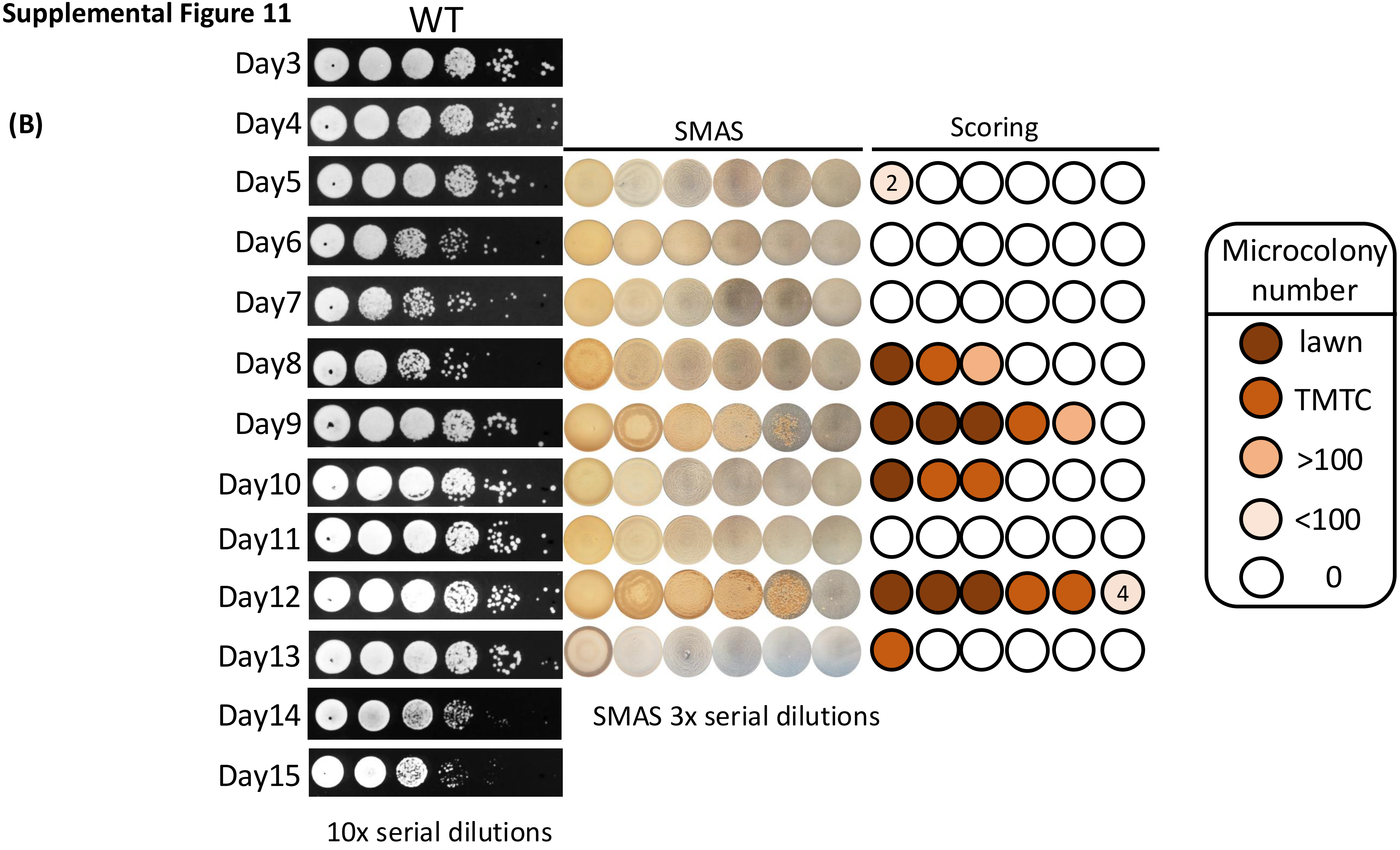

**Supplemental Figure 11**

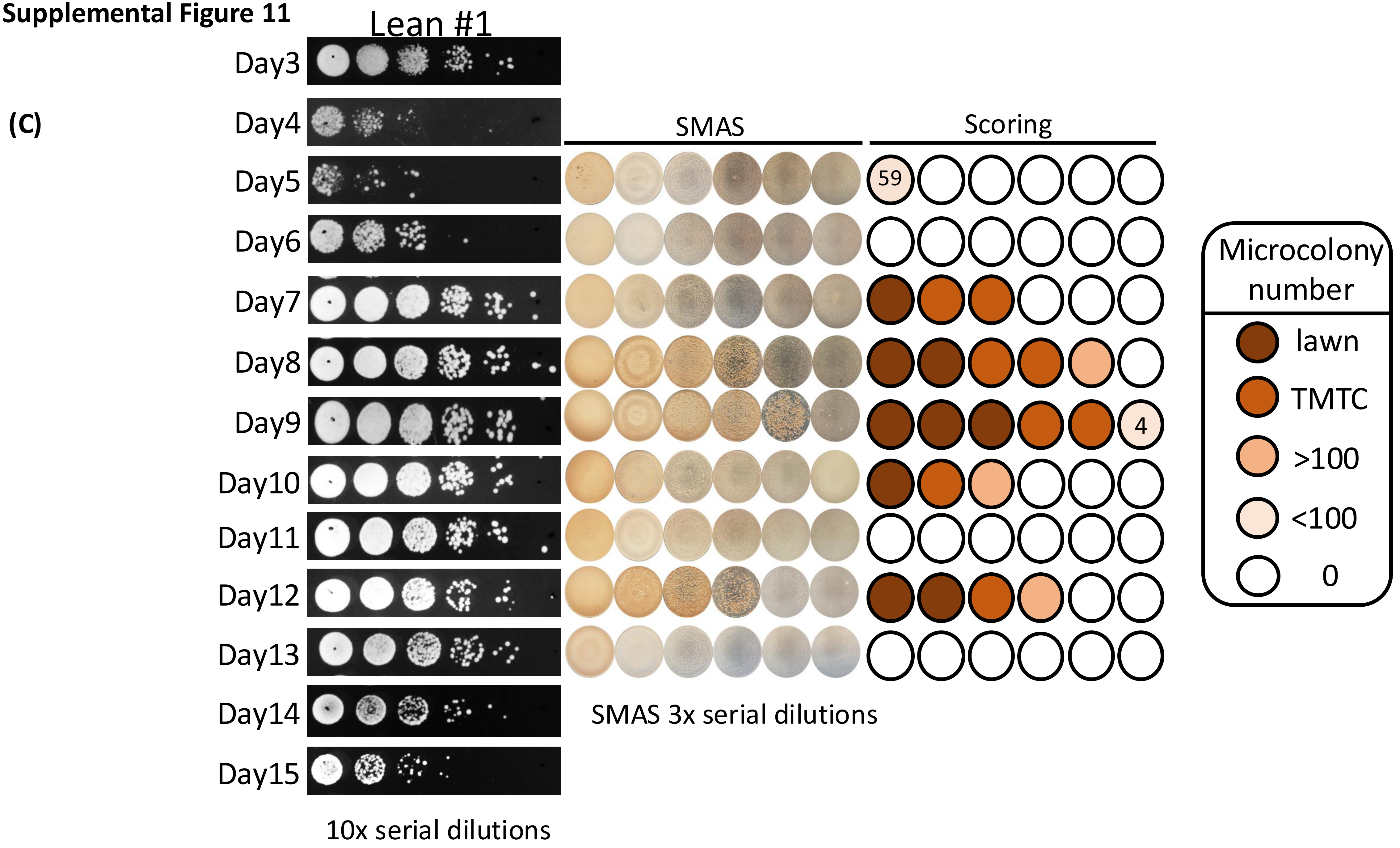

Supplemental Figure 11

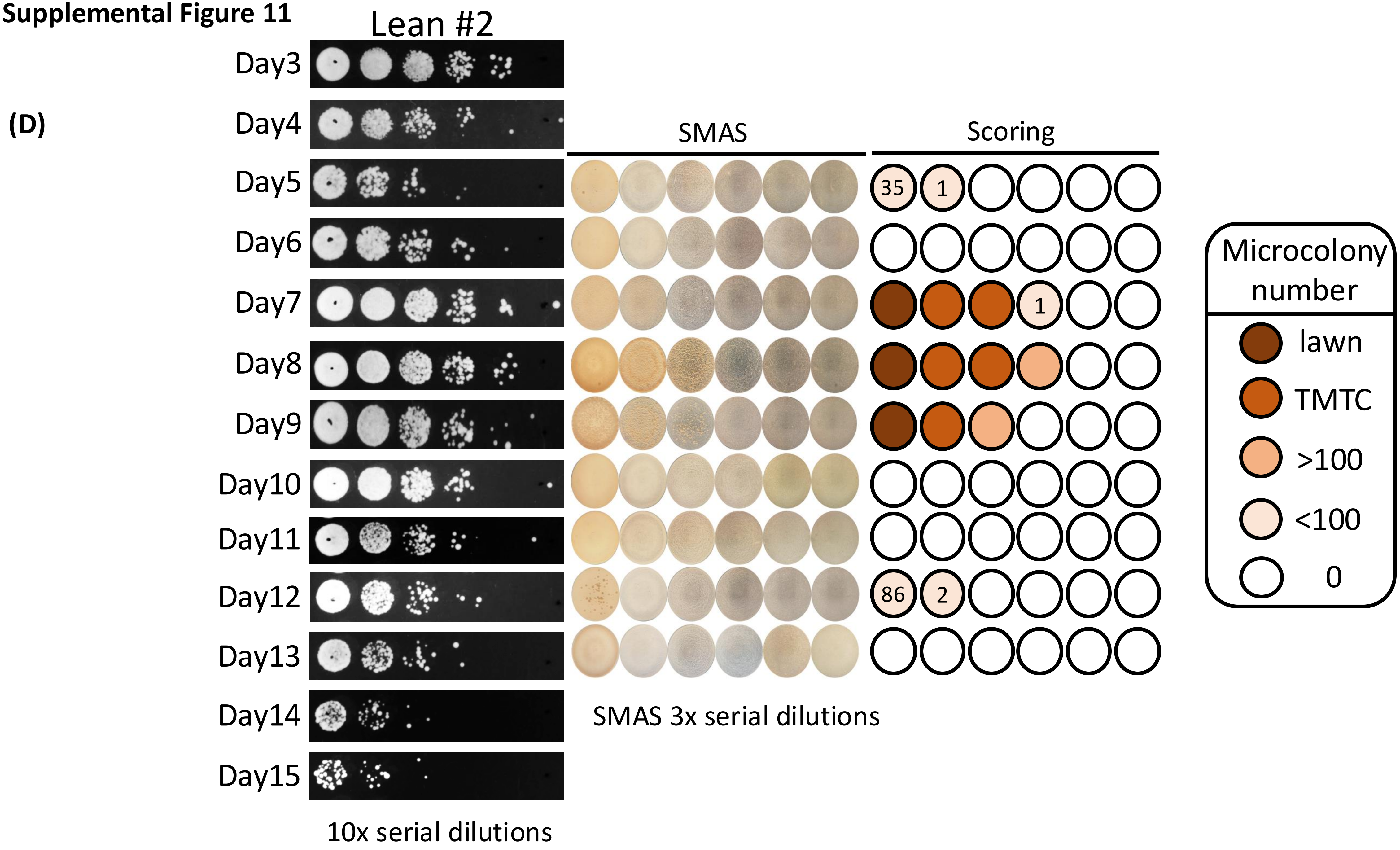

**Supplemental Figure 11**

(E)

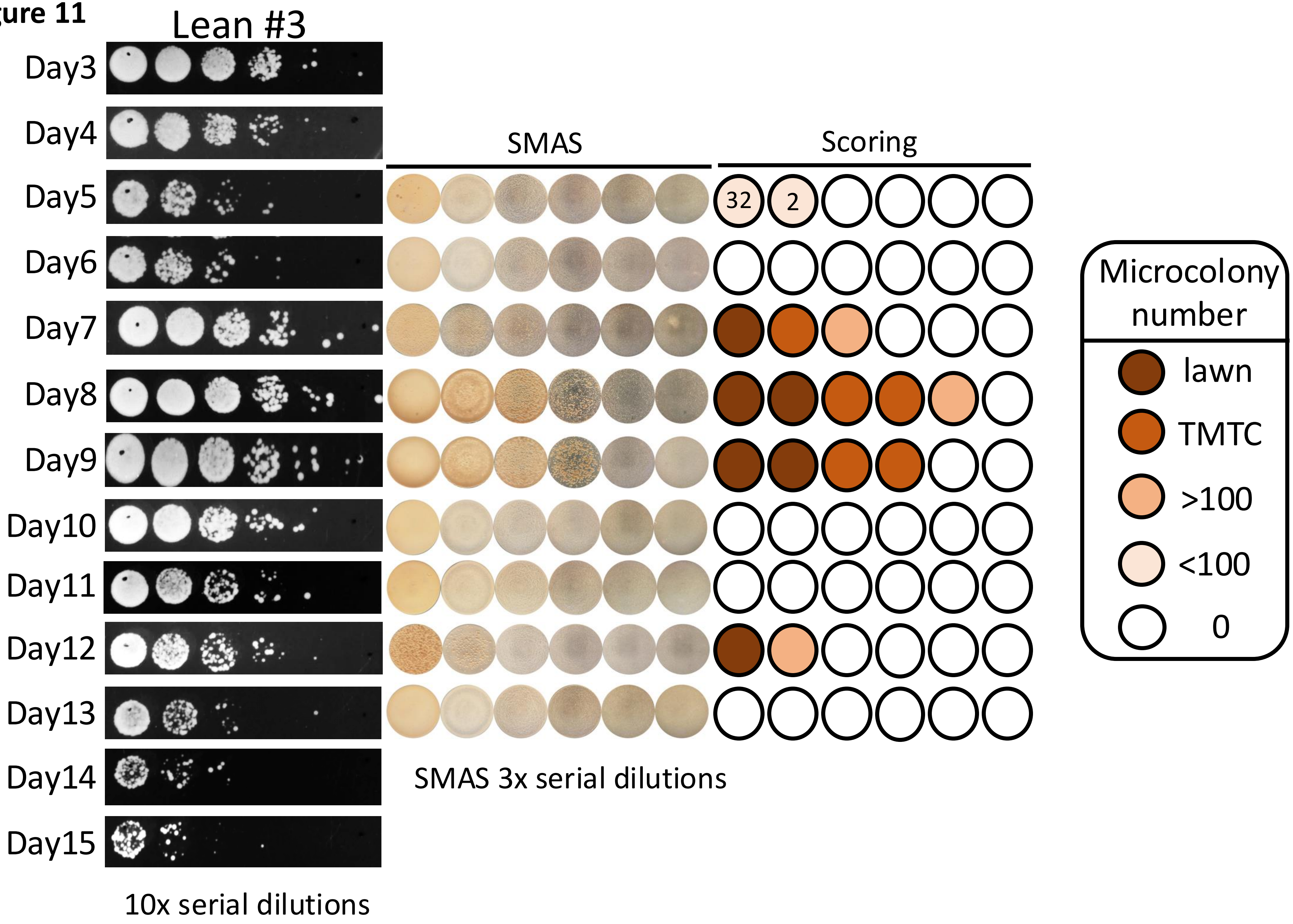

**Supplemental Figure. 12:**

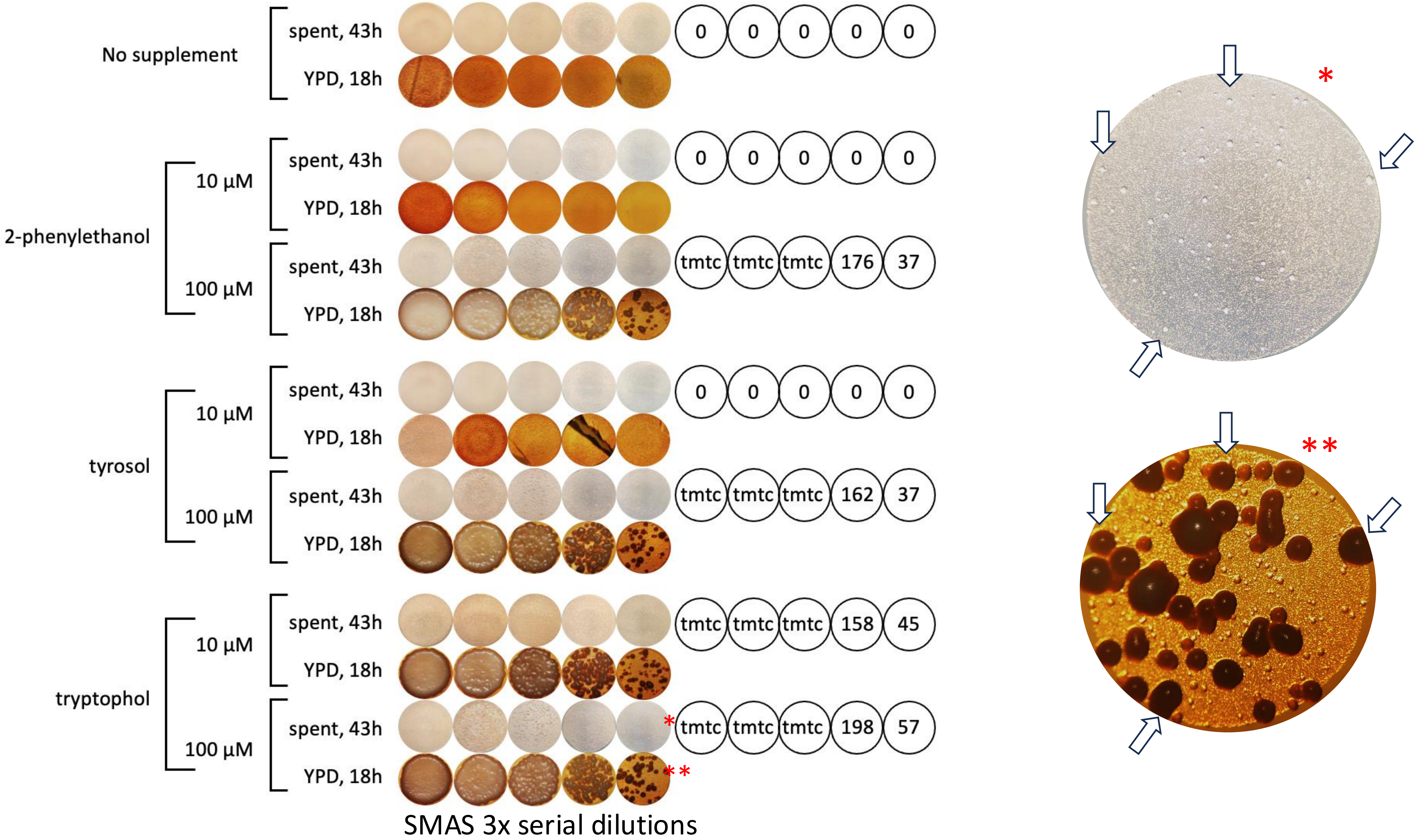

**Supplemental Figure 13.**

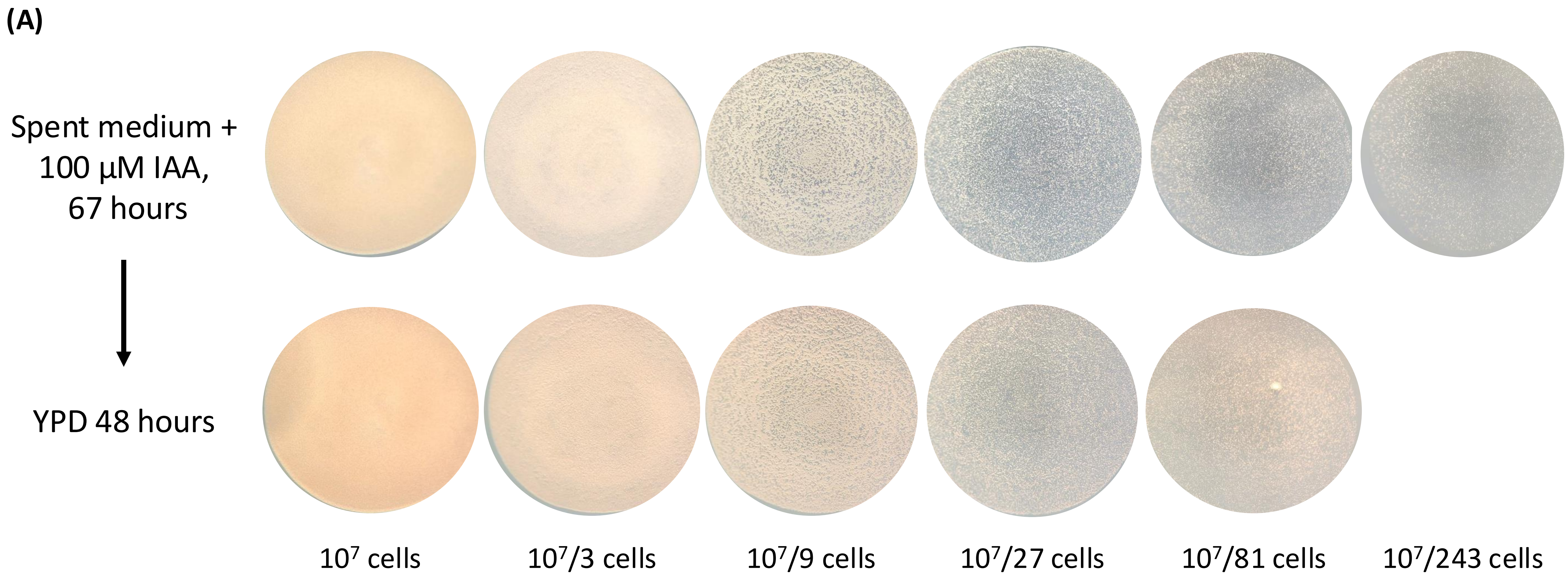

**Supplemental Figure 13.**

**(B)**

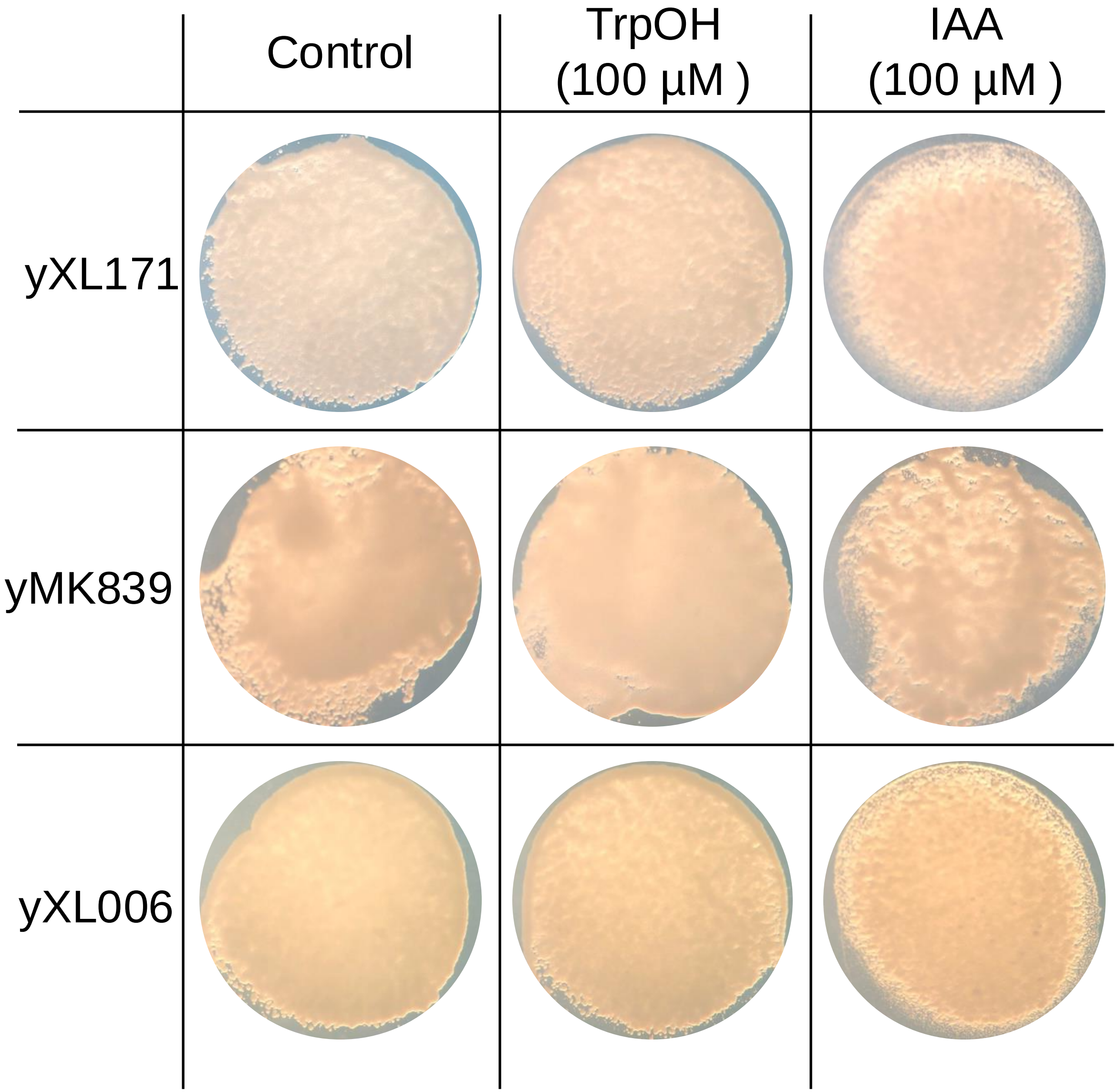

Supplemental Figure 14.

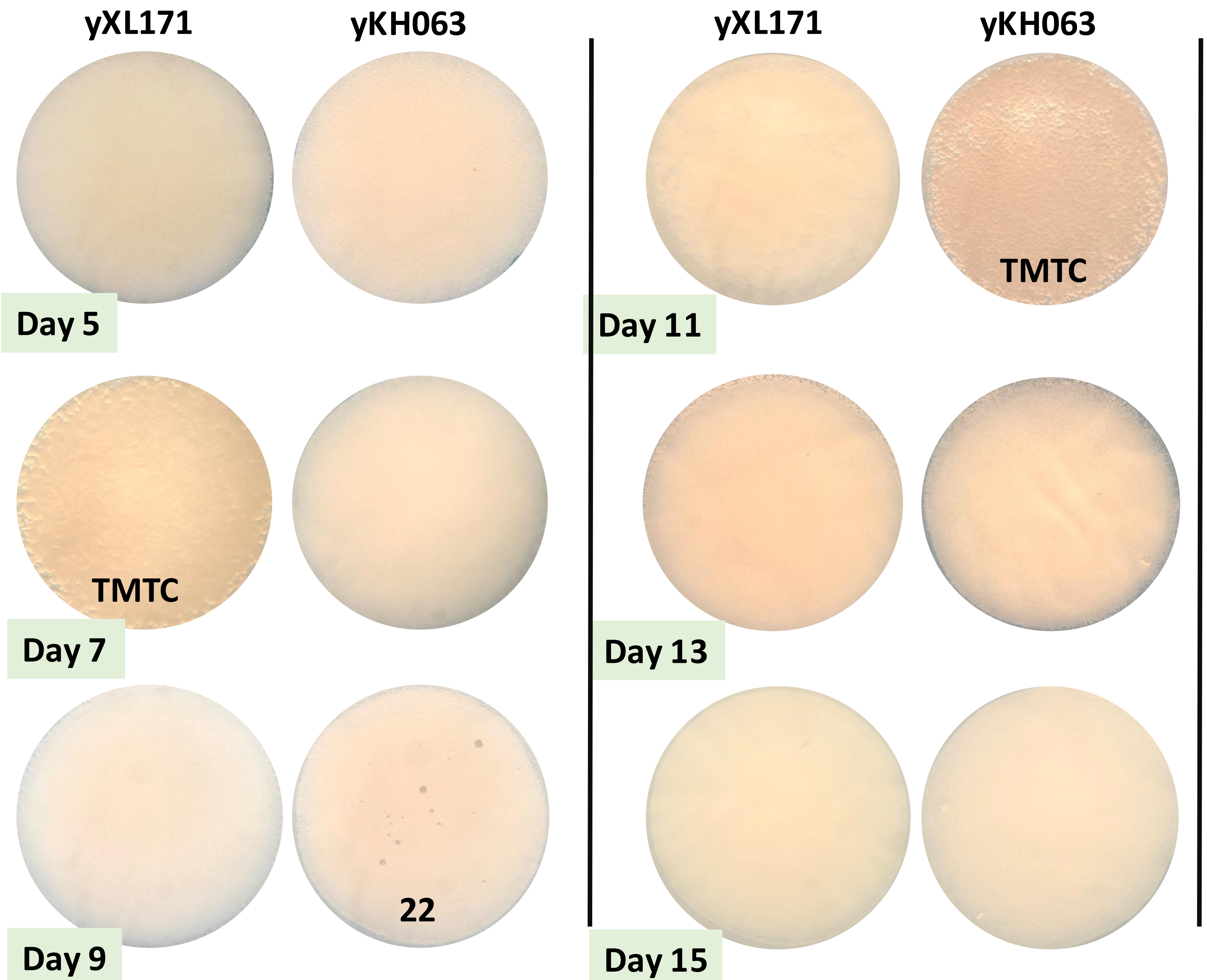


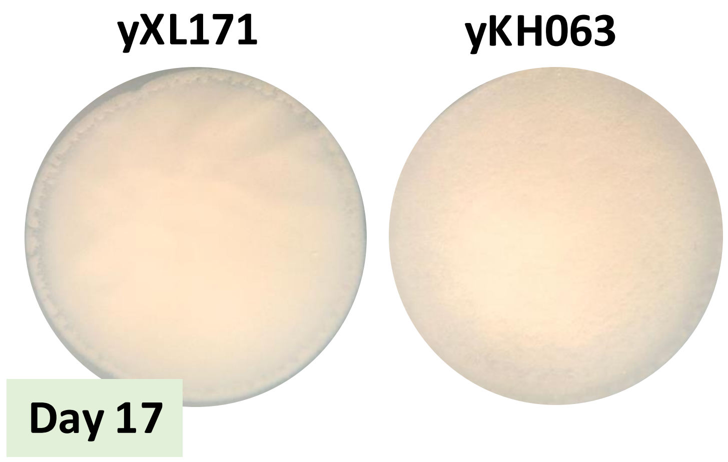


| Days | yXL171 (parental cell) | yKH063 *(aro8Δ; aro9Δ)* |
|---|---|---|
| 5 | 0 | 0 |
| 7 | TMTC | 0 |
| 9 | 0 | 22 |
| 11 | 0 | TMTC |
| 13 | 0 | 0 |
| 15 | 0 | 0 |
| 17 | 0 | 0 |

**Supplemental Figure 15.**

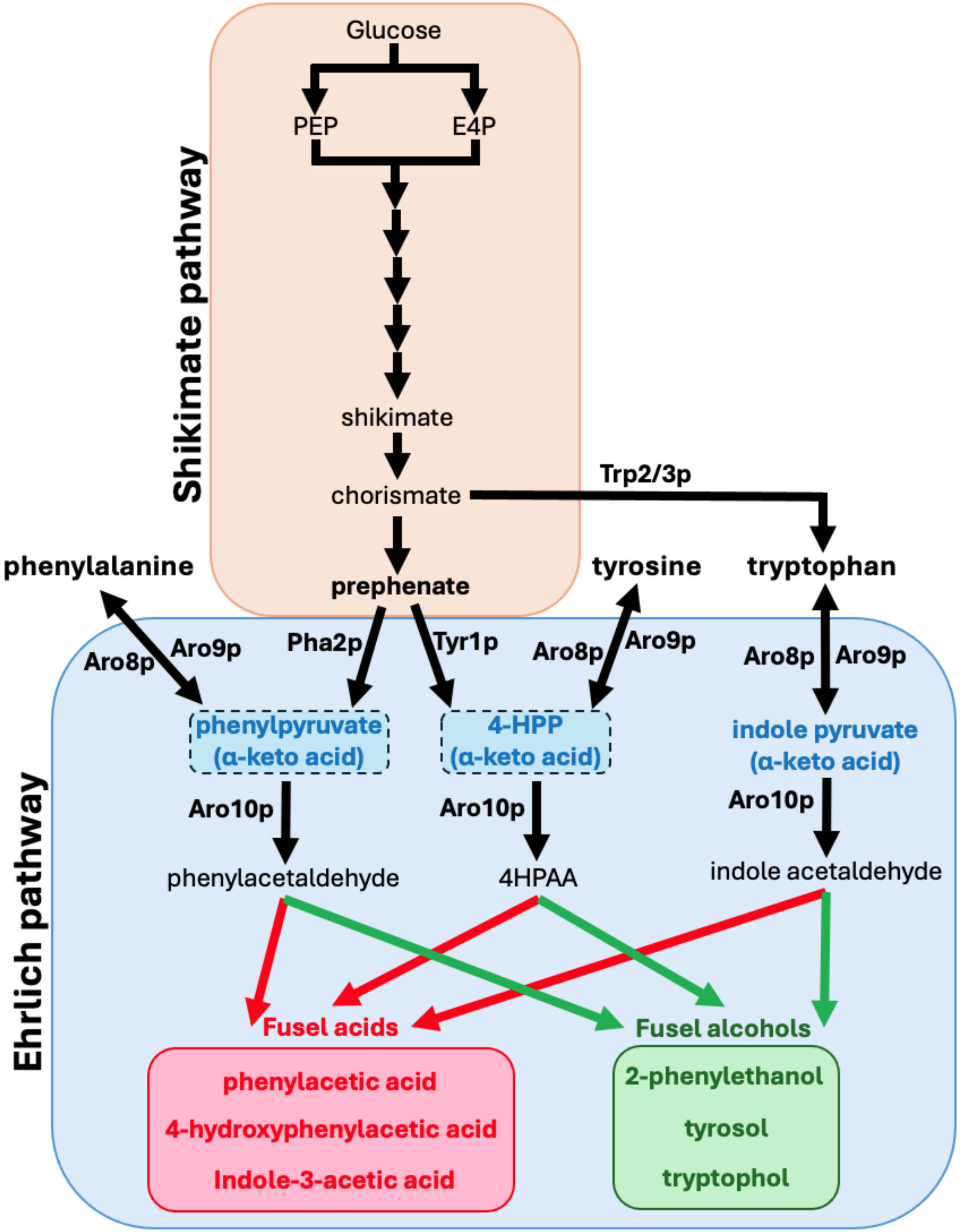
